# MOLECULAR DETERMINANTS OF THE ENDOCYTIC PROTEIN EPSIN CONTROLLING ITS LOCALIZATION AND FUNCTION IN CANCER CELL MIGRATION AND INVASION

**DOI:** 10.1101/2020.07.30.229195

**Authors:** Kayalvizhi Madhivanan, Lingyan Cao, Chris J. Staiger, R. Claudio Aguilar

## Abstract

Epsins are endocytic adaptor proteins with signaling and endocytic functions. The three mammalian epsin paralogs are made of an Epsin N-Terminal Homology (ENTH) domain and an unstructured C-terminal region. The highly conserved ENTH domain plays a role in signaling by blocking RhoGAP activity and is required for cell migration in mammalian cells. However, our lab has previously shown that only epsin full length overexpression can enhance cell migration, but the ENTH domain alone cannot. Among the three Epsin paralogs, epsin 3 followed by epsin 2 were able to substantially enhance cell migration. This study is the first one to systematically and comprehensibly address the contribution of different motifs within the epsin C-terminus to enhance protein localization and cell migration. We show that is not the lipid-binding ENTH domain, but the C-terminus of epsin the one playing a major role in epsin association with sites of endocytosis. Further, we dissected the contribution of individual C-terminal endocytic (clathrin-, AP2-, Ubiquitin- and EH domain-binding) motifs for epsin localization. We found that while all motifs show a degree of synergism, the clathrin-binding motifs are the most important for epsin localization. Our study also showed that, these motifs (particularly the clathrin binding site) play an important role in sustaining endocytic site dynamics and cell migration.

## INTRODUCTION

Epsins are a conserved family of endocytic adaptors characterized by an ENTH (<u>E</u>psin <u>N</u>-<u>T</u>erminal <u>H</u>omology) domain and an unstructured C-terminus (Fig.1). The ENTH domain of the epsins can bind to Phosphatidyl Inositol (4,5) bisphosphate (PIP_2_) at the plasma membrane and can induce membrane curvature. The C-terminus bears motifs for interaction with ubiquitinated cargo (*via* the Ubiquitin Interacting Motifs-UIM), EH domains (*via* Asn Pro Phe: NPF repeats), clathrin (Clathrin Binding Motifs-CBM), and AP2 (Aspartic acid, Proline, Tryptophan: DPW repeats) (Sen et al., 2012). Epsins can either function as endocytic accessory proteins, by binding to membranes and elements of the endocytic machinery to stabilize the endocytic interaction network, or they can function as adaptors, wherein they bind membrane, clathrin, and ubiquitinylated cargo proteins such as EGFR (Epidermal Growth Factor Receptor) and VEGFR2 (Vascular Endothelial Growth Factor 2) (Bertelsen et al., 2011, Kazazic et al., 2009, Sigismund et al., 2005, Pasula et al., 2012). This endocytic function of epsin is required for the signaling modulation of EGFR, VEGFR2 and the Notch signaling pathway (Wang et al., 2006a; Tian et al., 2004; Musse et al., 2012).

**Fig. 1.**
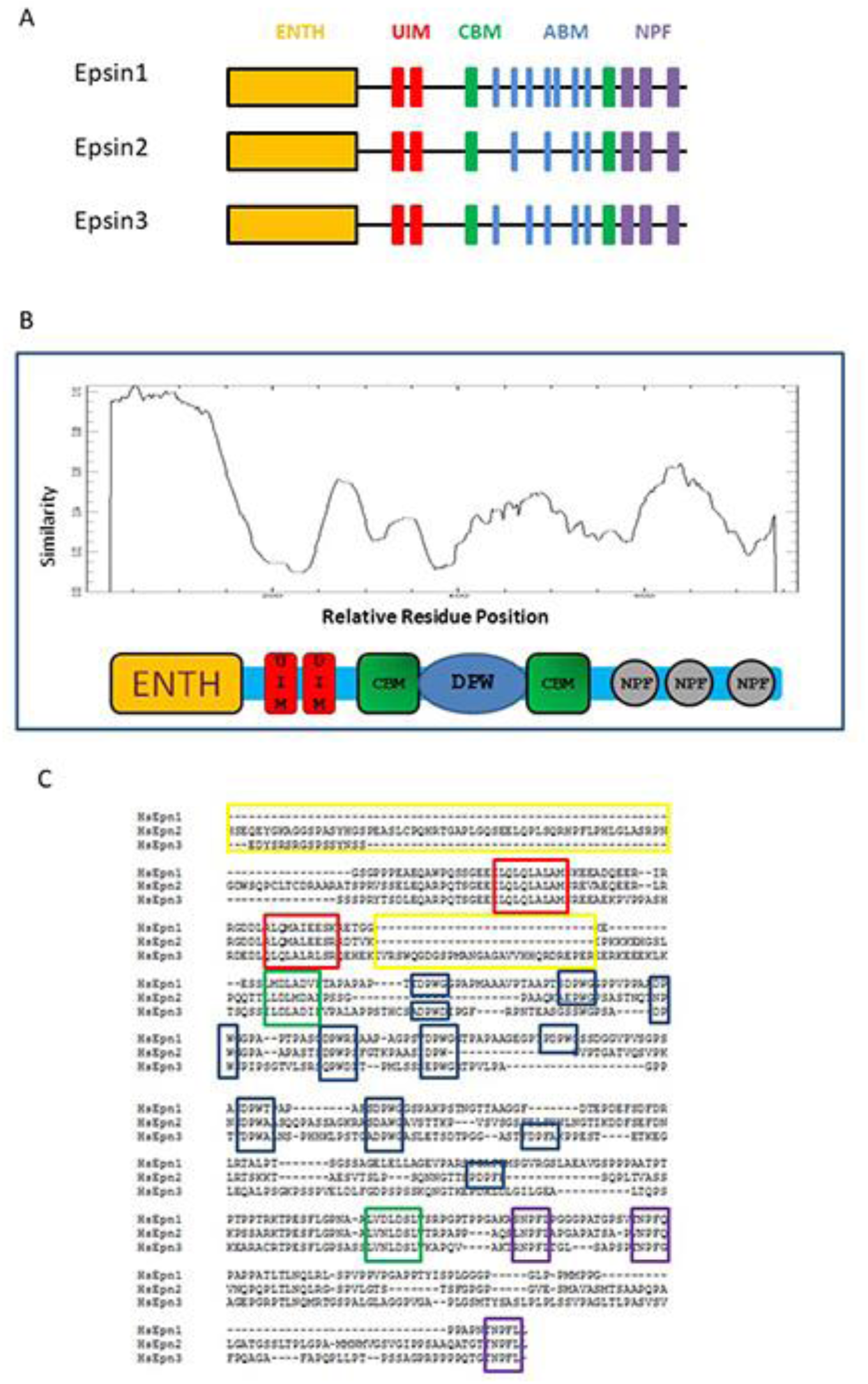
Three Mammalian Epsins Paralogs A) Domain architecture of the 3 mammalian epsins. B) Similarity graph for the 3 epsins shows the percentage conservation between the ENTH domains is very high. C) Sequence alignment of the C-terminus of the three epsins highlighting Ubiquitin interacting Motifs (UIMs) in red, Clathrin Binding Motifs (CBMs) in green, AP2 Binding Motifs (ABMs) (DPW/DPFs) in blue, NPFs (Asparagine, Proline, Phenylalanine) in purple and the paralog specific sequences in yellow. Additional likely AP2 binding domains have been highlighted (EPW/NPW/QPW).

Although other proteins within the endocytic network could potentially substitute for loss of epsins, these proteins are essential for embryo development and cell viability in eukaryotes (Fischer et al., 1997; Tian et al., 2004; Chen et al., 2009; Wendland et al., 1999, Aguilar et al., 2006). Yeasts have two epsins, Ent1 and Ent2. Simultaneous deletion of Ent1 and Ent2 is lethal; but viability is restored on expressing the ENTH domain of either epsin. It has been reported that the essential role of the ENTH domain lies in its signaling function, which depends on its physical interaction with Cdc42 GTPase Activating Proteins (Cdc42 GAPs: Bem3, Rga1 and Rga2) (Aguilar et al., 2006). The ENTH domain inhibits the function of Cdc42 GAPs and leads to the accumulation of activated, signaling competent, Cdc42-GTP. This signaling function was also conserved in mammalian epsins, where the mammalian epsin ENTH domain interacted with the mammalian RhoGAP RalBP1 and this activity was required for induction of cell migration and invasion (Coon et al., 2010). While the presence of an intact ENTH domain was essential for migration/invasion in epsin-depleted cells, enhancement of these processes only occur upon upregulation of full length epsins (Coon et al., 2010). This suggested a role for the epsin C-terminus in enhancing cell migration and invasion.

Sequence alignment of the three mammalian epsins shows that their ENTH domains are highly conserved, while the C-terminal sequences are the most divergent. In addition to the four kinds of endocytic determinants which are common to the C-terminus of the epsin paralogs, we identified paralog-specific sequences, which could account for their differential ability to enhance cell migration. The goal of the present study was to determine the role of the C-terminus in supporting the ENTH domain’s function in endocytic site dynamics and cell migration.

Endocytosis and membrane turnover at the leading edge is crucial for cell migration (Kural et al., 2015). Epsin has been shown to play critical roles in endocytosis as a checkpoint protein; and therefore, contributing to the proportion of abortive versus productive clathrin coated pits (Loerke et al., 2009). It is well established fact that epsin localizes to endocytic sites, however the major localization determinants are not known. This study shows that the C-terminus of epsin is the major determinant of localization. Further, we characterized the hierarchical requirement of the endocytic determinants found in the epsin C-terminus for localization. Based on these findings, we were able to propose a model for epsin recruitment to endocytic sites. Additionally, we also were able to address fine differences between the three epsin paralogs by comprehensive mutational analysis.

## RESULTS

### Epsin C-terminus if the major determinant of localization

Epsin localizes to the endocytic sites as marked by clathrin and AP2 at the plasma membrane. While the ENTH domain of epsin binds to PI(4,5)P_2_ enriched in the plasma membrane, the C-terminus (Ct) binds to different components of the endocytic machinery. However, it is not known which the major localization determinant of epsin is. To resolve this question we chose epsin 2 because it shared commonalities with the other 2 paralogs. Epsin 1 and 2 are ubiquitously expressed, while epsin 2 and 3 overexpression enhanced cell migration and invasion (Sen et al., 2012). We rationalized that based on the results from epsin 2, key mutants could be chosen and tested from epsin1 and 3.

We transfected full length (FL) epsin2-GFP, ENTH2-GFP and Ct2-GFP (del ENTH2) and EV (empty vector – GFP alone) in Hela cells and fixed the cells to observe if the ENTH domain or Ct of epsin by itself could localize to endocytic sites. The ability to localize at endocytic sites was assessed by their co-localization with AP2. AP2 and clathrin being the major markers of clathrin-mediated endocytic (CME) sites, we chose AP2 because it is found only at the plasma membrane, while clathrin has been observed on endosomes and also at the trans-golgi network (TGN) (Keyel et al., 2004). In addition, we measured the mislocalization of these constructs in other cellular compartments, namely the TGN, nucleus and the cytoplasm. FL-epsin2 occupied almost every endocytic site as marked by AP2, showing co-localization of 98% (Fig.2). While ENTH2 did not localize at endocytic sites, the Ct2 showed a high co-localization of 96% with AP2. In addition to localization at the endocytic sites, Ct2 also mislocalized to the cytoplasm and TGN (Fig.2). This showed that while the C-terminus of epsin is the major determinant of localization, the ENTH domain supports the C-terminus. This property of the epsin C-terminus was the same across all the paralogs. We speculated that the TGN localization of the Ct2 could be due to the epsin CBMs binding to clathrin at the TGN in the absence of the ENTH domain which kept epsin targeted to the PIP_2_ enriched plasma membrane (unlike its paralog EpsinR which is found in the TGN).

**Fig. Error! No text of specified style in document.**
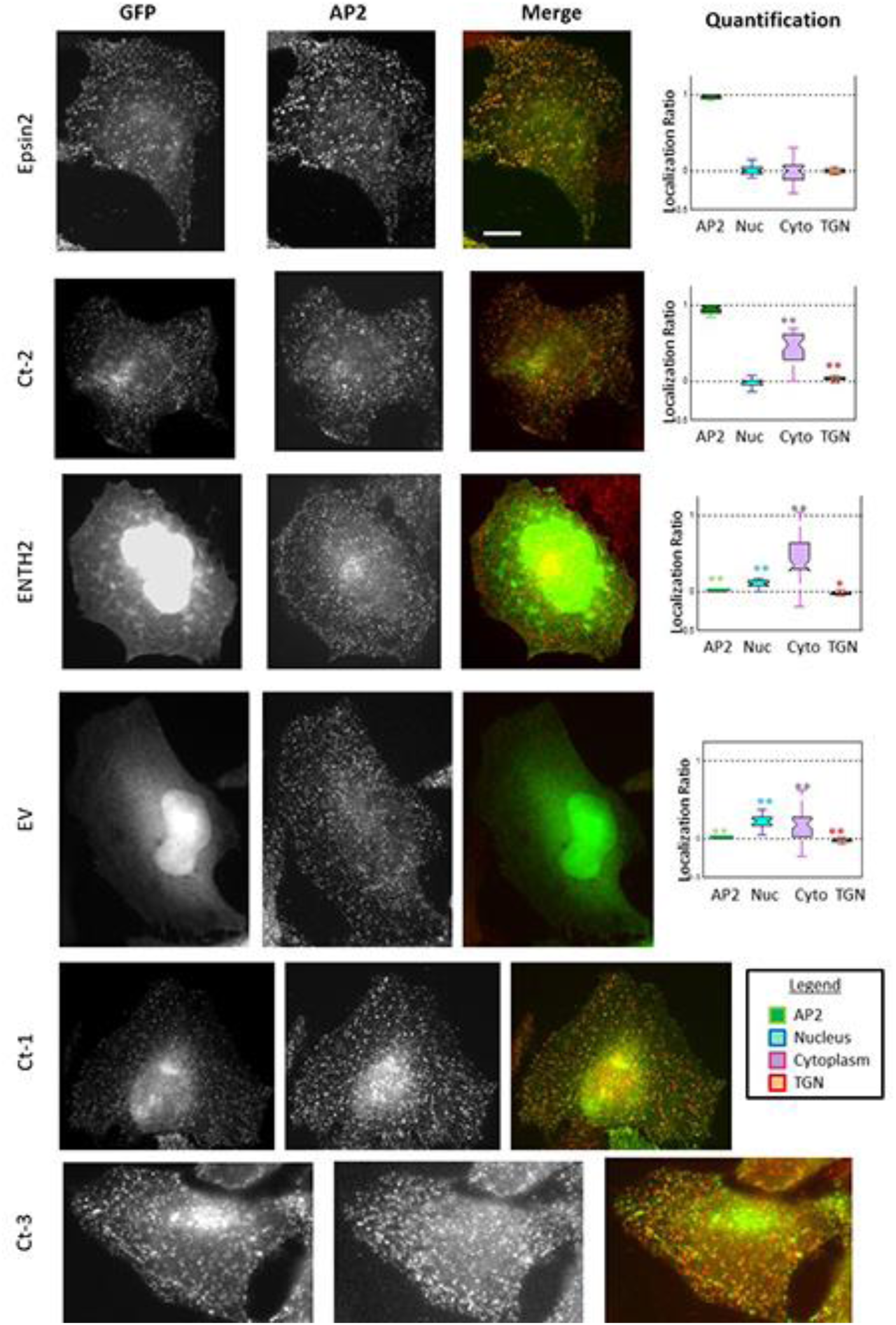
Determination of co-localization of ENTH versus the C terminus of Epsin paralogs. Representative images show co-localization of GFP empty vector (EV), GFP-epsin 2-GFP, GFP-ENTH2, GFP-Ct2 (C terminus of epsin2), Ct1 (C terminus of epsin1) and Ct3(C terminus of epsin3) with AP2. Comprehensive quantitative analysis done for epsin 2 is shown in the right most panels, depicting co-localization with AP2, cytoplasmic, TGN and nuclear staining. The values were normalized against epsin 2 full-length. ** signifies p<0.05 by Wilcoxon test. Scale Bar = 10uM

### The endocytic determinants within the C-terminus contribute to localization

The C-terminus of epsin has defined regions which can bind to four different components of the endocytic machinery, as described before. These endocytic motifs only contribute to 10% of the C-terminus sequence (Fig.2). The next step was to determine the contribution of various C-terminal determinants to epsin localization. We started with the common determinants within the epsin Ct which interact with the endocytic machinery and prepared mutants using a systematic and comprehensive approach. We performed combinatorial mutagenesis in FL and Ct constructs of epsin2 (Fig. 3). Epsin2 has two ubiquitin interacting motifs (UIM), two clathrin binding motifs (CBM), seven AP2 binding motifs (DPW) and 3 EH-domain binding motifs (NPF), will be referred to as UCDN2 (epsin2 WT). We started by mutating each motif individually and named the mutant following the motifs that were still intact, for e.g., U_2_CDN2 refers to epsin2 with the first UIM mutated; UC_2_DN2 refers to epsin2 with CBM1 mutated and so on. Following this, we made 4 constructs with all the endocytic motifs of one kind mutated where CDN2 would be epsin2 with both UIMs mutated; UCN2 would be epsin2 with all the 7 DPWs mutated and so on. For full list of constructs and their description used in this study, see Materials and Methods. To test co-operativity between the endocytic motifs, we made all possible combinatorial mutants from CDN2, UDN2, UCN2 and UCD2 constructs as shown in Figure 3. The next row of mutants have 2 types of endocytic determinants mutated (DN, CN, CD, UN, UD, UC) and the following has 3 types of endocytic determinants mutated (U, C, D, N). The final mutant ucdn has all the endocytic determinants mutated. All these constructs were also obtained in the context of the Ct of epsin2 alone (ENTH domain deleted).

**Fig. 1.**
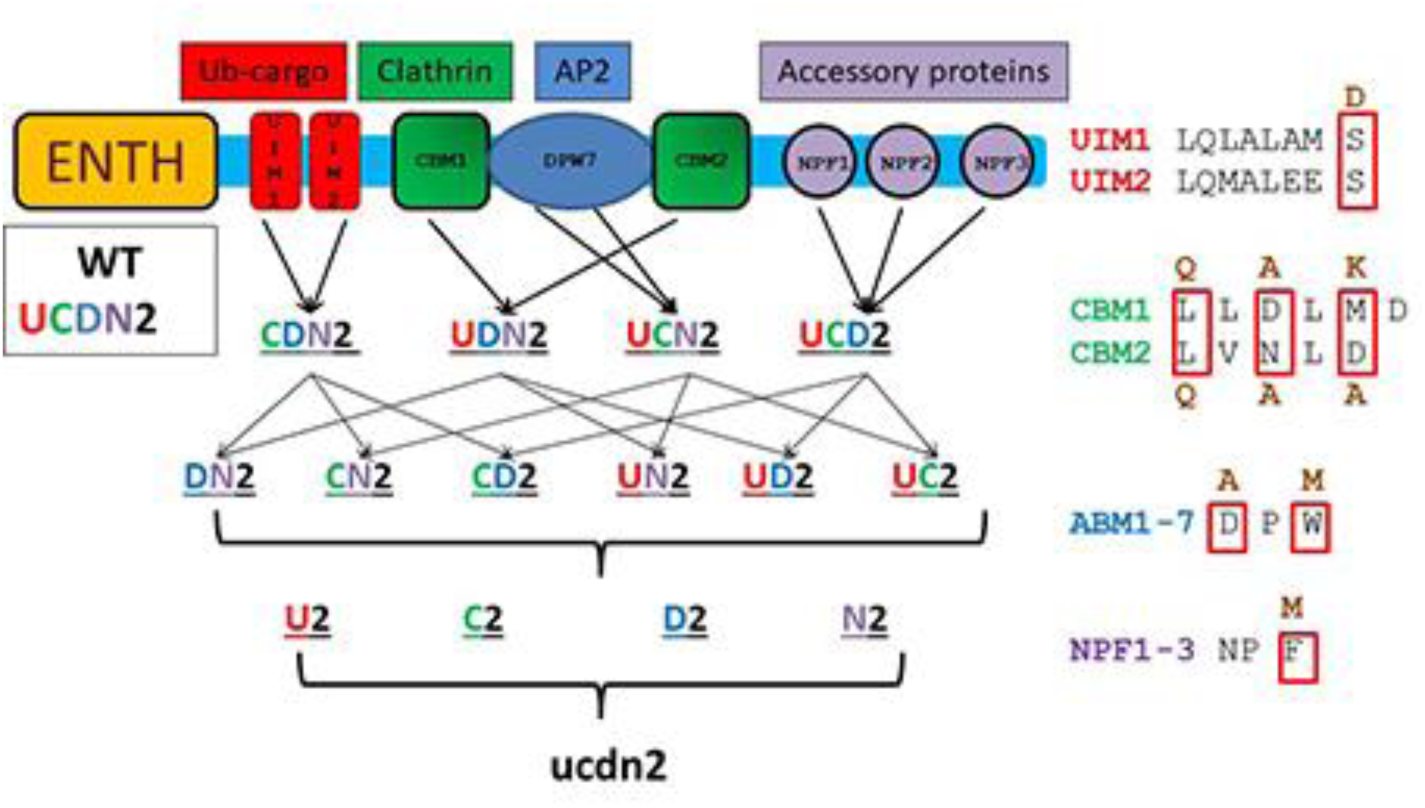
Combinatorial mutations made in epsin 2. UCDN refers to the WT sequence. The first row (CDN, UDN, UCN and UCD) refers to mutants with one type of endocytic determinant mutated. The second row (DN, CN, CD, UN, UD, UC) represent mutants with two types of endocytic determinants mutated. Third row (U, C, D, N) refers to mutants with three types of endocytic determinants mutated. The WT motif sequence along with the mutations introduced are shown on the right. Please refer to text for details.

In order to determine if the endocytic determinants contribute to localization we began the analysis with ucdn2 and Ct-ucdn2. Mutation of the 4 types of endocytic determinants in the context of the FL epsin2 could not form puncta and exhibited a complete loss of co-localization with AP2, showing a localization pattern similar to the EV (Fig.4). Both the mutants had diffused cytoplasmic staining and nuclear staining. Previously, we observed that Ct2 had gained TGN, which was significantly lost after mutation of the endocytic determinants (Ct-ucdn2). The above results suggested that the endocytic determinants, which contribute to only 10% of the C-terminal sequence, are the major determinants of epsin2 localization at endocytic sites.

**Fig. 2.**
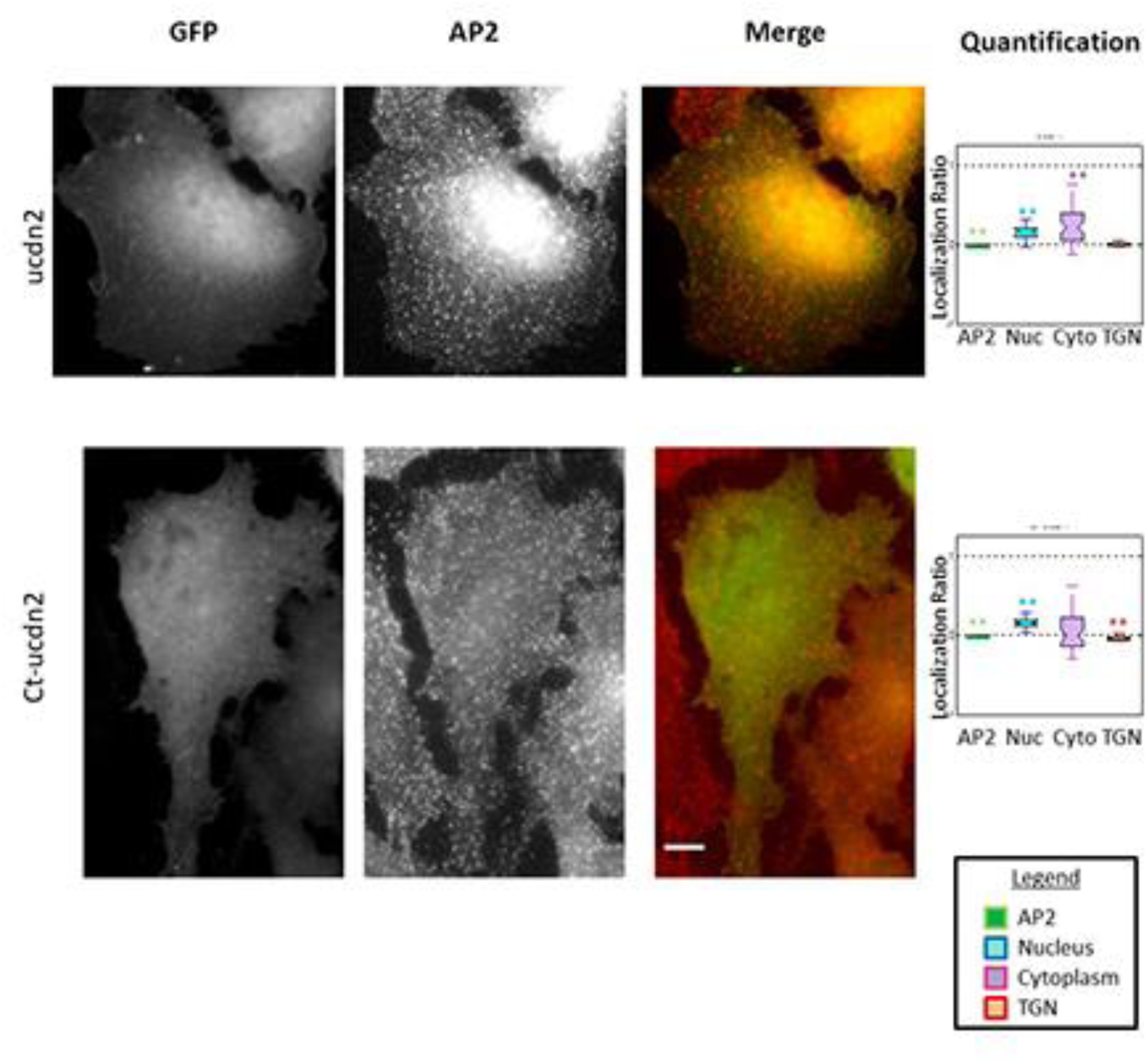
The endocytic determinants in the Ct of epsin are the major determinants of localization. Representative images and quantification of the mutant lacking intact endocytic determinants in the context of the FL and Ct-epsin 2 are shown here along with the quantification. The data is normalized with respect to FL and Ct-epsin 2 respectively for ucdn2 and Ct-ucdn2. ** signifies p<0.05 by Wilcoxon test. Scale Bar = 10um

### Clathrin Binding Motifs and AP2 Binding motifs (ABM) are the most important localization determinants

Since the mutants ucdn2 and Ct-ucdn2 were unable to localize at endocytic sites, we started our co-localization analysis from mutants which contained only 1 kind of endocytic determinant intact. None of the mutants from this group could localize at endocytic sites, even in the presence of the ENTH domain (Fig.5). An interesting observation was that C2 (which had both CBMs intact) was the only mutant to be excluded from the nucleus, and also showed enrichment at the TGN, suggesting that it was non-specifically binding to clathrin at the TGN.

**Fig. 3.**
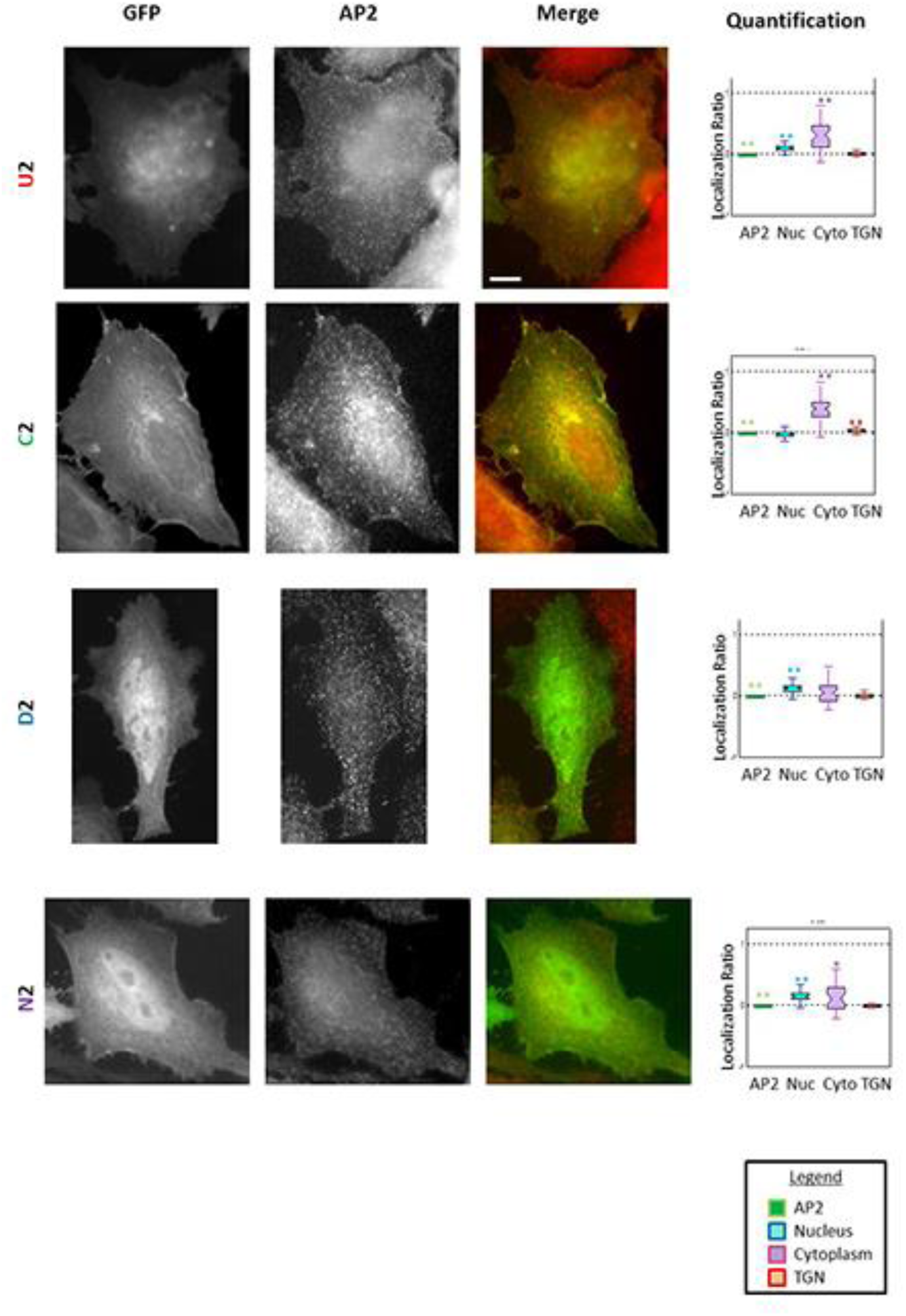
Localization of epsin2 mutants with only 1 kind of endocytic determinant intact. Representative images of the mutants are shown here along with their quantification normalized wrt FL epsin 2. ** signifies p<0.05 by Wilcoxon test. Scale Bar = 10um

Analysis of the next group of mutants, which comprises of six mutants with two kinds of endocytic motifs intact, gave interesting results wherein the mutations in the context of the FL versus the Ct were different. The analysis of the 6 mutants in the absence of the ENTH domain resulted in 2 groups with different localization patterns. One group which consisted of Ct-UC2, Ct-CD2 and Ct-CN2 had the ability to localize at endocytic sites showing high co-localization with AP2 (Fig.6 A). Another group consisting of Ct-UD2, Ct-UN2 and Ct-DN2 was unable to show punctuate localization (Fig.6 B). The common determinant lacking in the latter group was the absence of the CBMs, which clearly suggests that the CBMs are the most important determinant of localization. Consistently, mutants of the latter group (lacking functional CBMs) showed nuclear staining and a significant decrease in TGN staining as compared to Ct2. However, these same mutations introduced in FL epsin2, showed that only UN2 was unable to localize at endocytic sites (Fig.7). The mutant UN2 lacks intact CBMs and ABMs which suggested that the CBMs and ABMs are the two most important determinants of localization. The localization pattern of UD2 and DN2 in a FL versus Ct-2 backbone shows punctuate and diffused localization respectively further reinforces the role played by the ENTH domain in supporting the C-terminus for proper epsin2 localization (Fig.6 B, Fig.7).

**Fig. 4.**
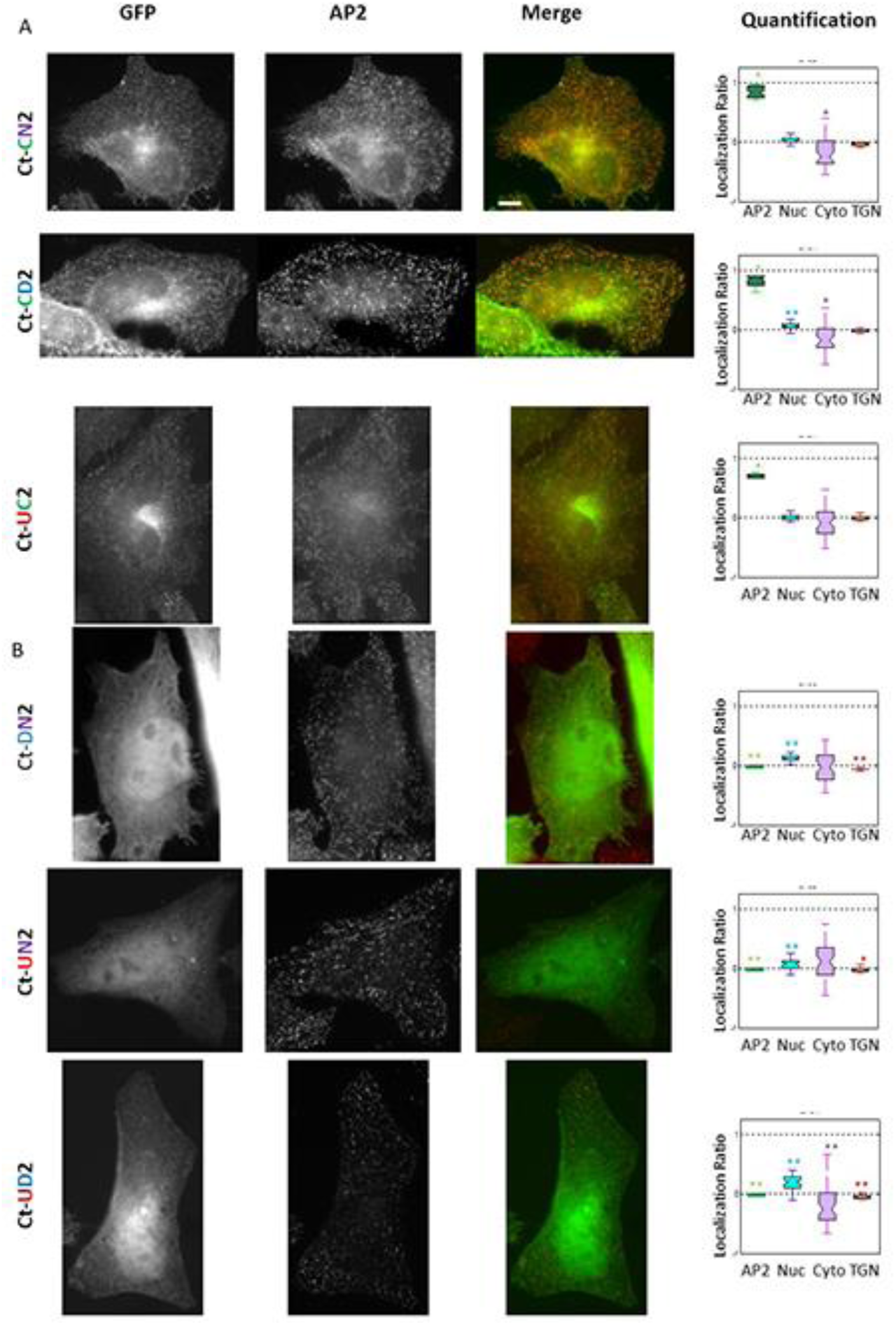
CBM is the strongest localization determinant. Representative images and quantification of the mutants with 2 kinds of endocytic determinants in the context of Ct-2 are shown here along with the quantification normalized with respect to Ct-2. A) These mutants have intact CBMs along with one other kind of endocytic determinant intact. B) These mutants lack CBMs along with another kind of endocytic determinant. ** signifies p<0.05 by Wilcoxon test. Scale Bar = 10um

**Fig. 5.**
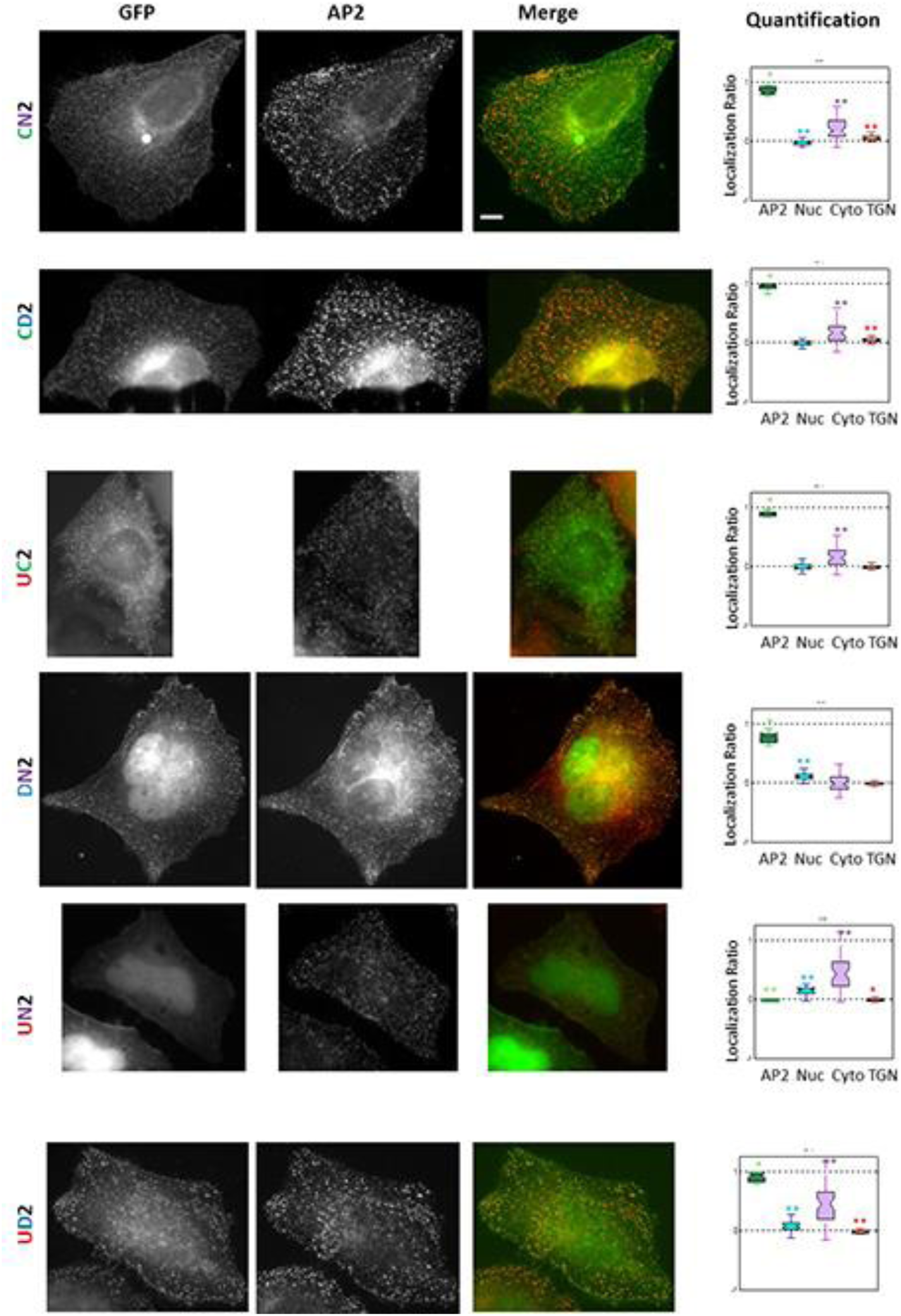
CBM and DPws are the strongest localization determinants. Representative images and quantification of the mutants with 2 kinds of endocytic determinants in the context of FL epn2 are shown here along with the quantification normalized with respect to FL epn2. A) These mutants have intact CBMs along with one other kind of endocytic determinant intact. B) These mutants lack CBMs along with another kind of endocytic determinant. ** signifies p<0.05 by Wilcoxon test. Scale Bar = 10um

Next, we analyzed the mutants which had only one kind of endocytic determinant mutated and the other 3 intact. None of the mutants namely, CDN2, UDN2, UCN2 and UCD2 showed a significant decrease in co-localization with AP2 (Fig.8), even in the absence of the ENTH domain (Fig.9). Gain in nuclear staining and loss of the TGN staining was again observed upon mutation of the CBMs. This suggested that there is redundancy and co-operativity among the different C-terminal determinants.

**Fig. 6.**
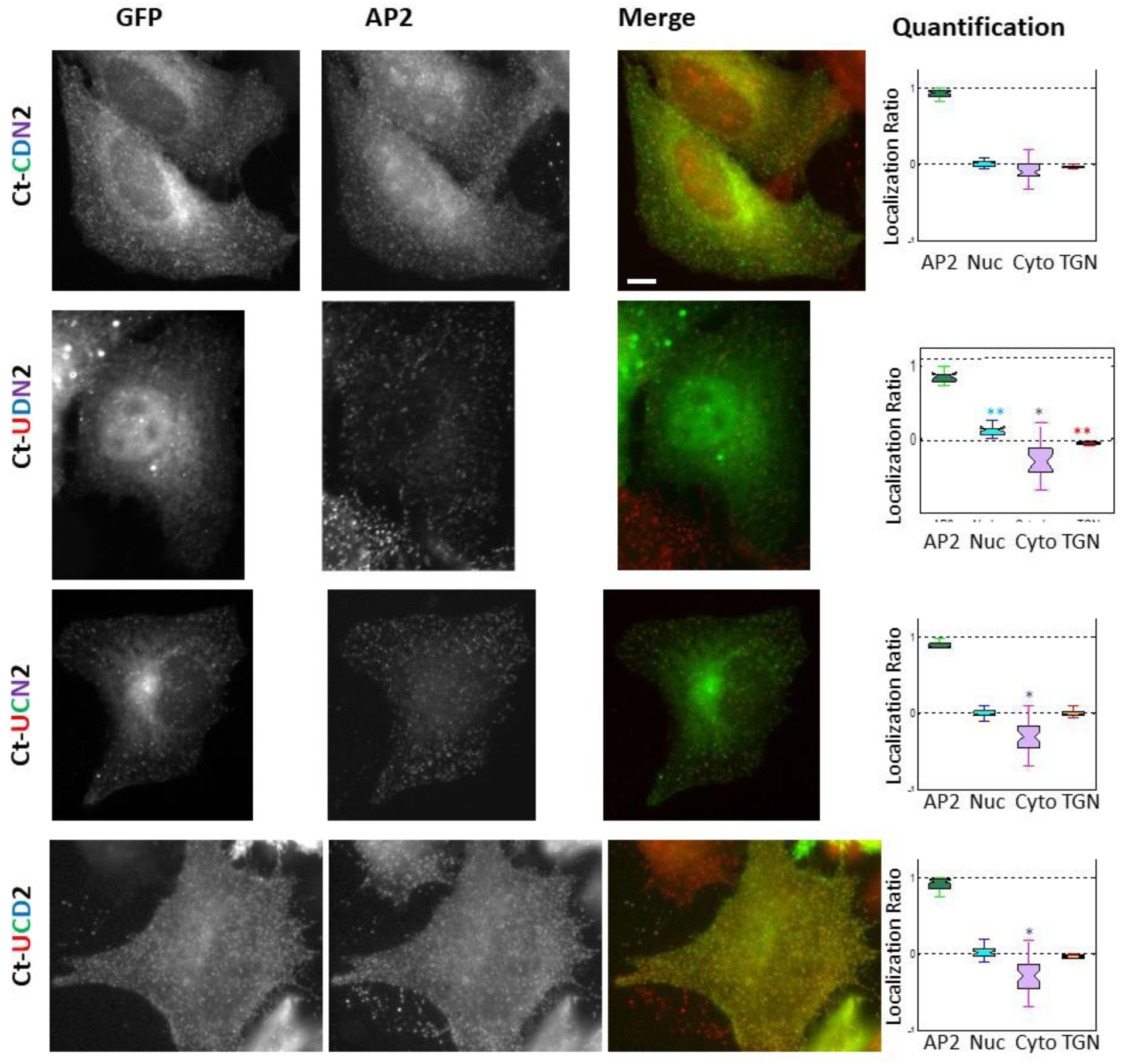
Redundancy and co-operation among the different endocytic determinants. Representative images and quantification of the mutants with 3 kinds of endocytic determinants in the context of Ct-2 are shown here along with the quantification normalized with respect to Ct-2. ** signifies p<0.05 by Wilcoxon test. Scale Bar = 10um

**Fig. 7.**
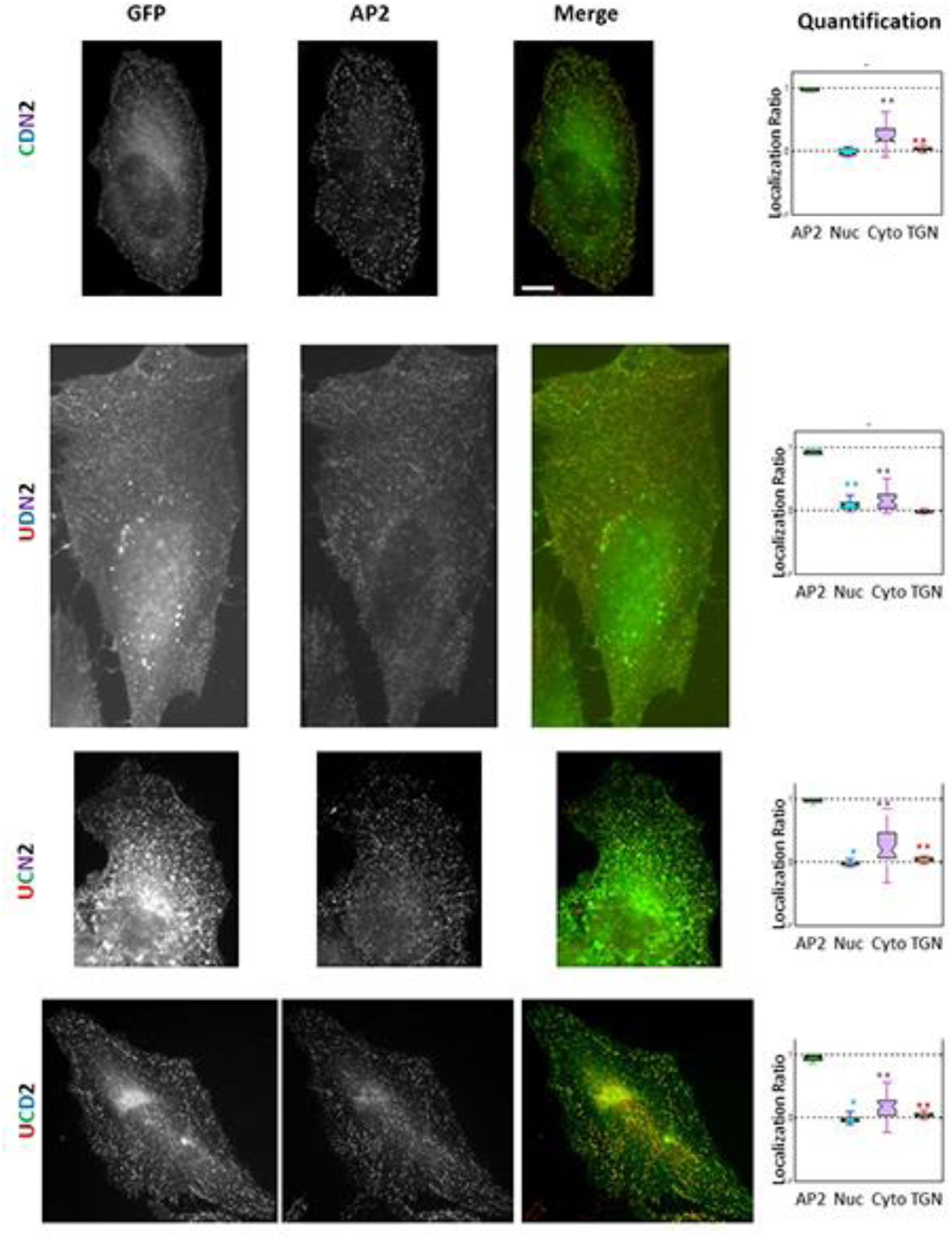
Redundancy and co-operation among the different endocytic determinants. Representative images and quantification of the mutants with 3 kinds of endocytic determinants in the context of FL epn2 are shown here along with the quantification normalized with respect to FL epn2. ** signifies p<0.05 by Wilcoxon test. Scale Bar = 10um

### CBM2 is more important than CBM1 for epsin2 localization

CBMs being the strongest localization determinant, we questioned if CBM1 or CBM2 was equally important for epsin2 localization. The 2 CBMs in epsin fall under 2 different classes. CBM2 has a more classical clathrin box motif (LФpФp, where Ф represents hydrophobic residue and p represents polar residue) LVNLD, while CBM1 has a different consensus LLDLMD (Drake et al., 2000). When we mutated CBM1 or CBM2 separately in the context of the full length construct, we did not see any significant difference in localization patterns. So, we took advantage of our combinatorial mutants to establish the relative importance of the CBMs. The mutant Ct-UN2 (CBMs and DPWs mutated and lacking the ENTH domain) shows no localization at endocytic sites. So, we reintroduced either intact CBM1 or CBM2 in this mutant and analyzed its localization at endocytic sites. While Ct-C1UN2 showed only 30% co-localization with AP2, Ct-C2UN2 showed 70% co-localization with AP2 (Fig.10 A). This finding agreed with previous results where clathrin binding to the distal CBM2 was higher than CBM1 (Drake and Traub, 2001). Further, introducing an intact CBM2 to replace CBM1 (CS1) or CBM1 to replace CBM2 (CS2) by themselves (Fig.10 B) or along with their flanking regions (CF1 and CF2 respectively) (Fig.10 C) in the Ct-UN2 mutant showed a localization pattern similar to the CtUN-2 mutant. Our experiments revealed that the distal CBM2 was more important for epsin2 localization to endocytic sites and so is its position along the C-terminus of epsin. It is noteworthy that in all 3 epsin paralogs, out of the two CBMs, the classical CBM is towards the distal end.

**Fig. 8.**
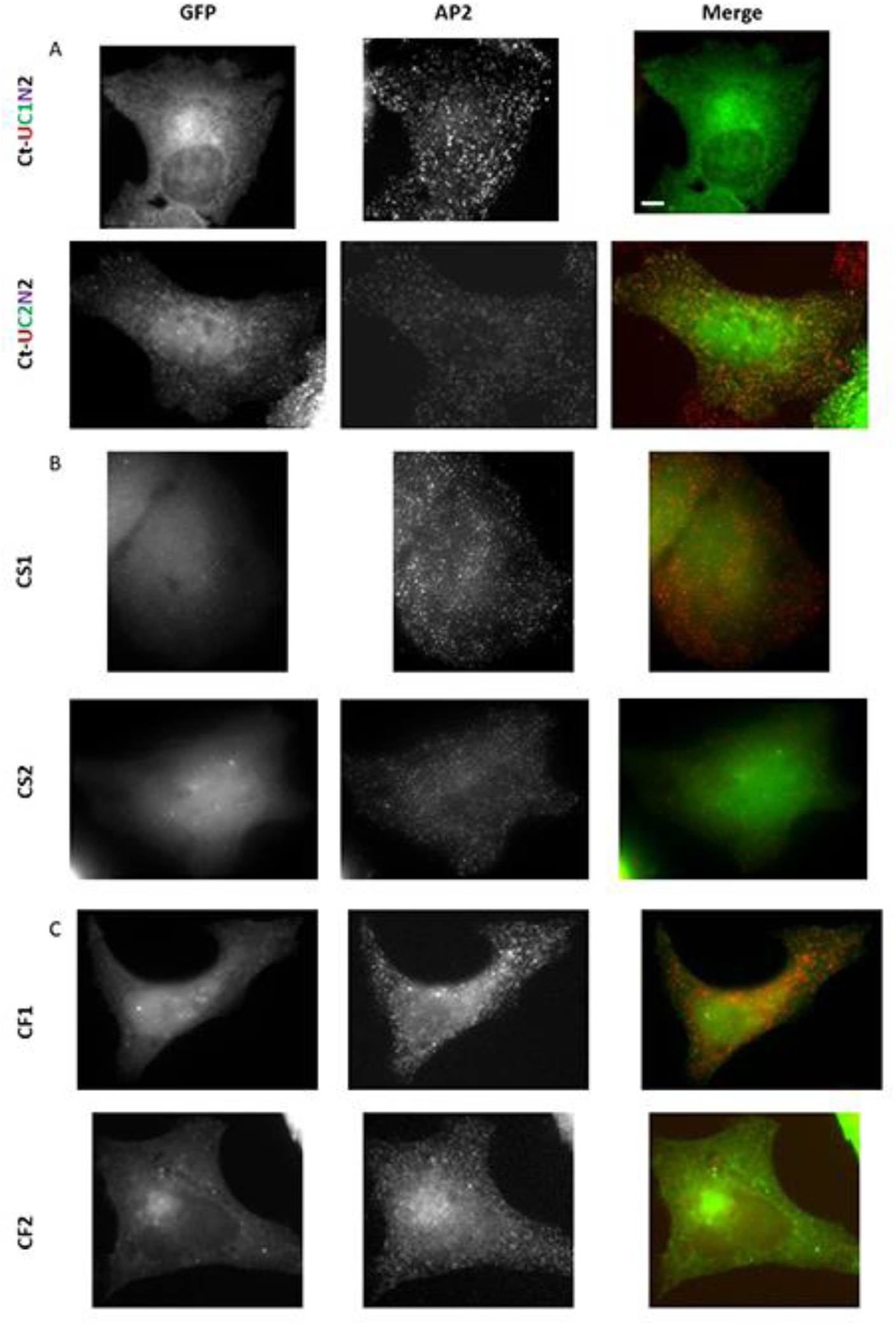
CBM2 is a stronger localization determinant than CBM1 for epn2 localization. The base mutant used in all these constructs is Ct-UN2. A) An intact CBM1 or CBM2 was introduced. The mutant Ct-UC2N2 rescued the localization defect of CtUN-2. B). CS1-An intact CBM2 was reintroduced in the position of CBM1. CS2-An intact CBM1 was reintroduced in the position of CBM2. Neither CS1 nor CS2 was able to rescue the localization defect of epn2. C) CF1-CBM2 and its respective flanking region replaced CBM1 and its flanking region. CF2 – CBM1 and its respective flanking region replaced CBM2 and its flanking region. Scale Bar = 10um

### Comparison of the hierarchy of the localization determinants

The three epsin paralogs have the 4 types of endocytic motifs and co-localize with each other and with AP2 almost to 100% (Fig.11). We further co-transfected FL epsin1 along with epsin2 mutants (with only 2 types of endocytic determinants intact). When epsin1 was expressed at higher levels, it was able to displace epsin2 mutants from endocytic sites (Fig.12). These results clearly displayed that the three epsin paralogs occupied the same endocytic sites. So, we expected that the hierarchy of the endocytic determinants would be the same among the three paralogs.

**Fig. 9.**
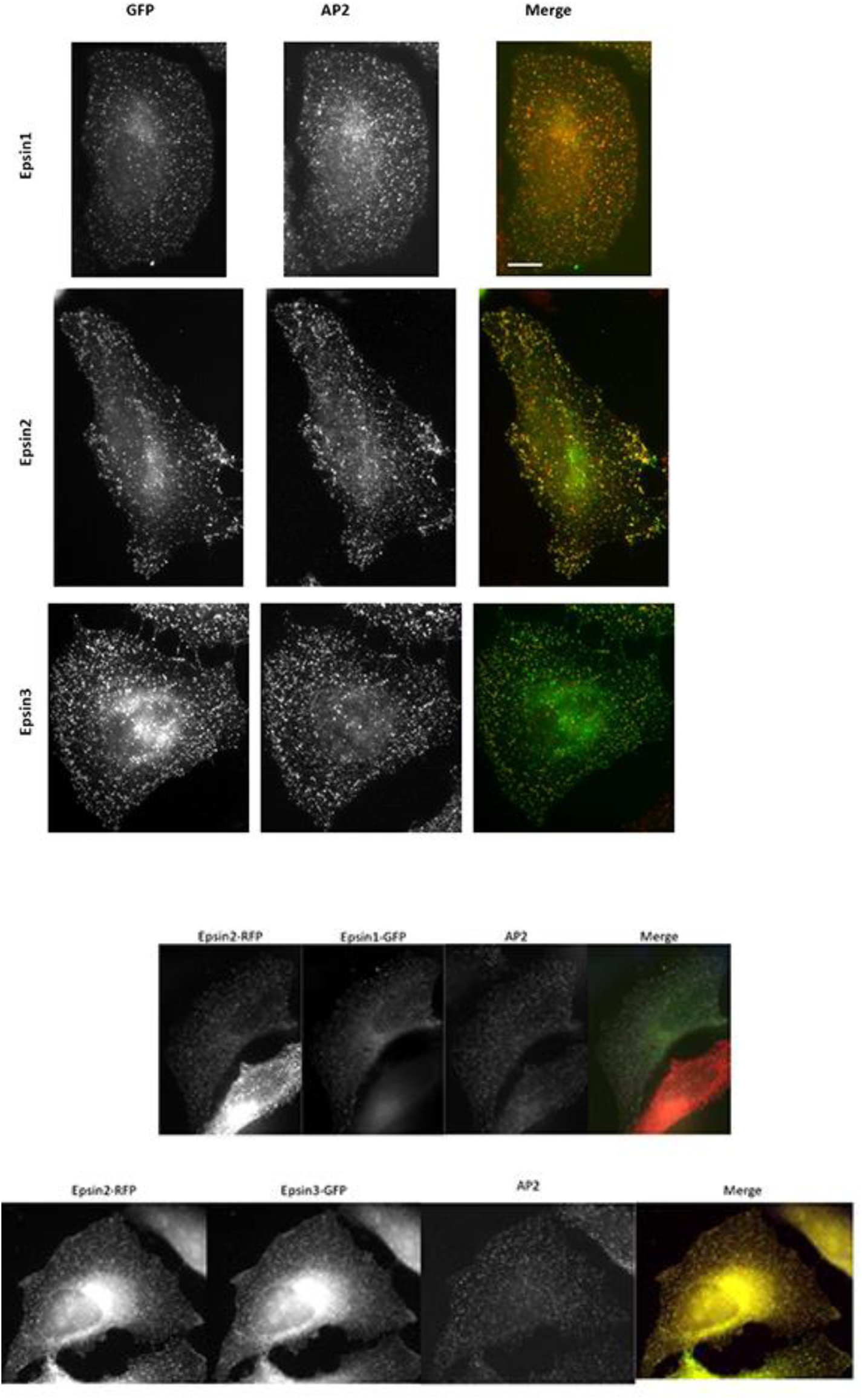
The co-localization of the three epsin paralogs with AP2. These images show that epsin1, 2 and 3 co-localize with AP2 at endocytic sites. Scale Bar = 10μm

**Fig. 10.**
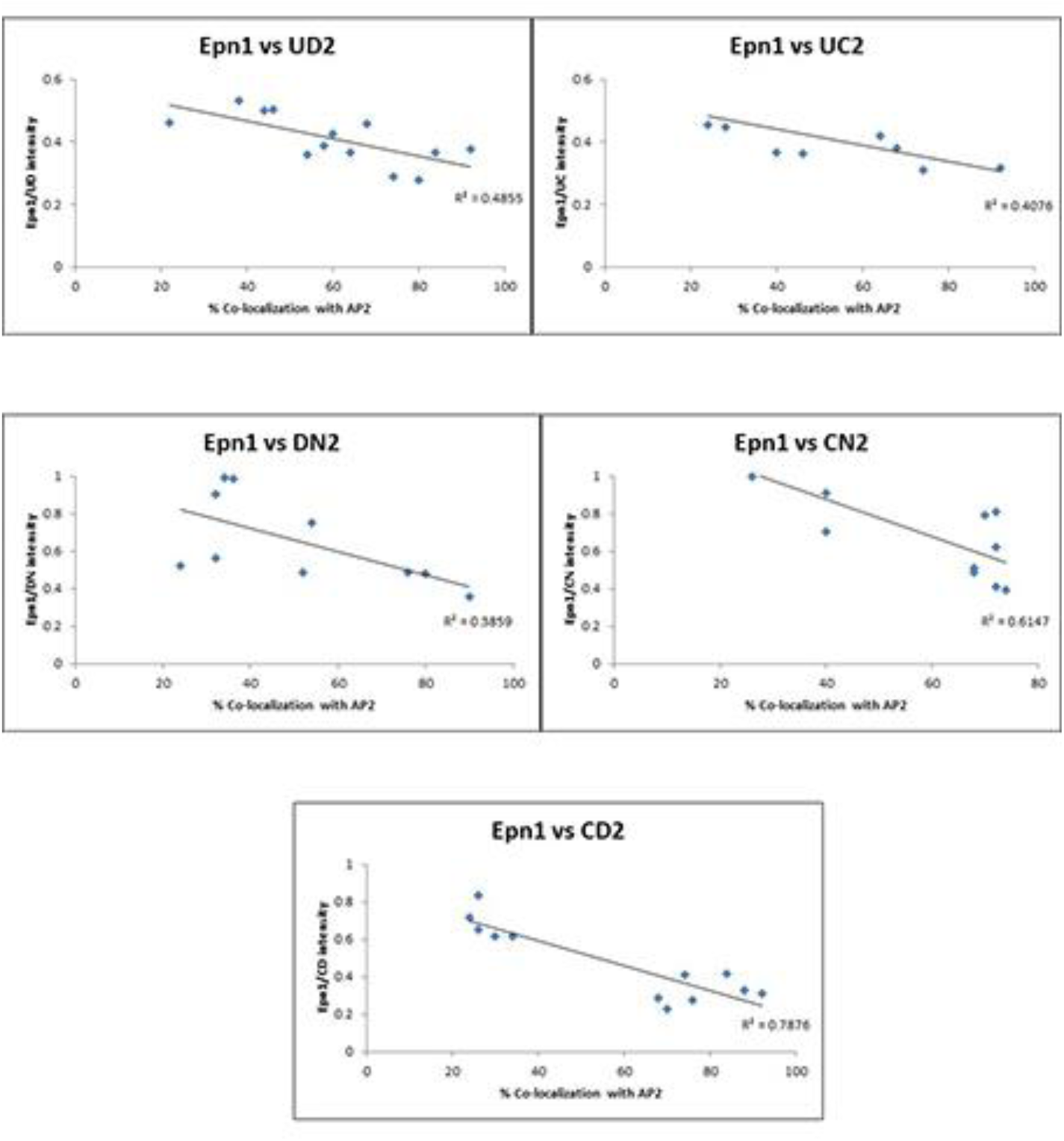
Displacement of epsin2 mutants by FL epsin1. The graphs show a correlation between Epsin 2 mutants co-localization with AP2 as a function of Epsin1 to Epsin2 mutant intensity ratio.

Mutations were introduced in epsin1 and epsin3 (Fig.13). We analyzed the mutants following the bottom-up approach as we did for epsin2. ucdn1 and ucdn3, like ucdn2, did not localize to endocytic sites (Fig.14). Next, we analyzed the mutants which have only 1 kind of endocytic determinant intact in FL epsin1 and 3 (Fig.15, Fig.16). This result was very similar to epsin2, where none of the mutants belonging to this group could localize to endocytic sites.

**Fig. 11.**
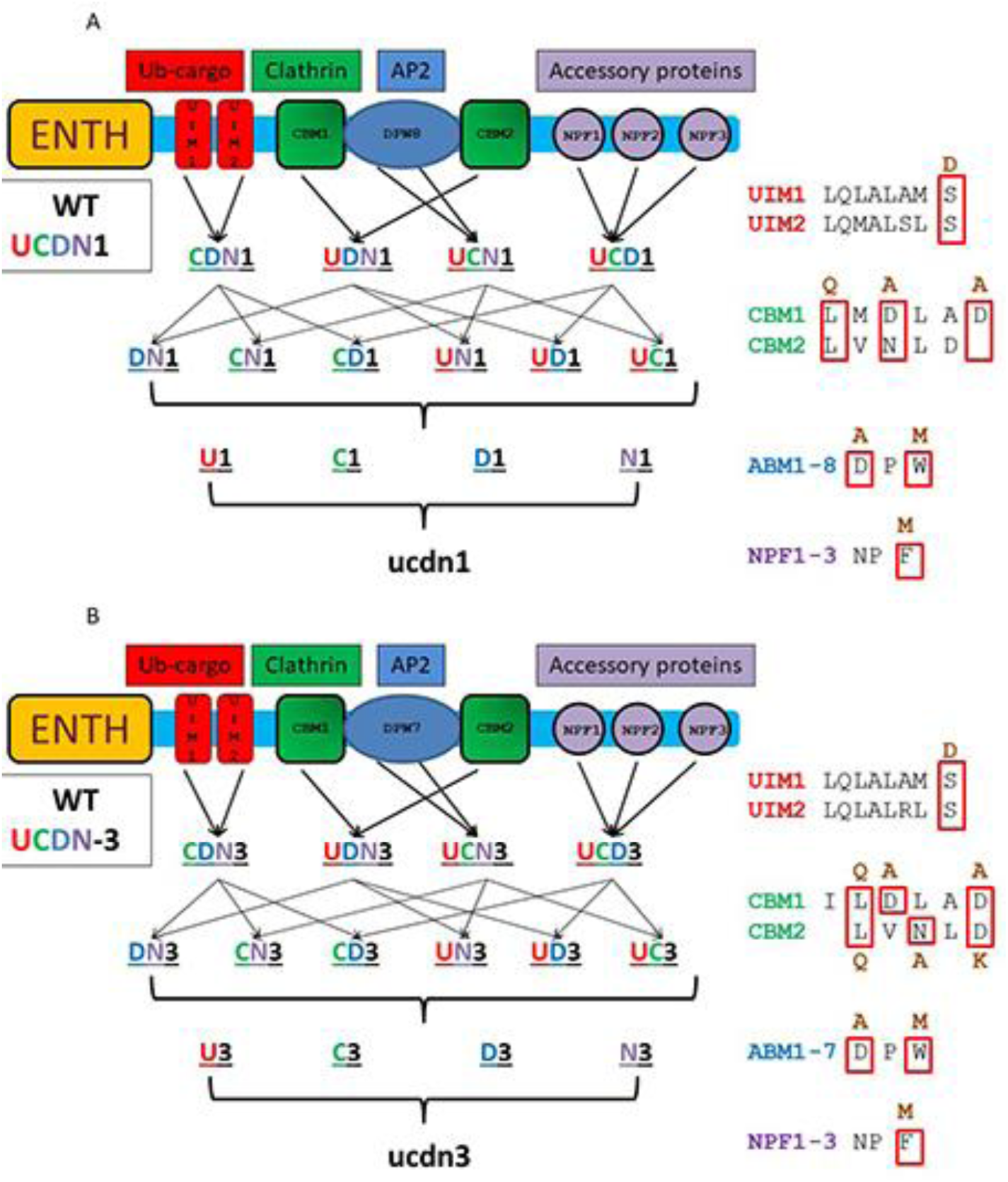
Combinatorial mutations made in epsin1 (A) and epsin3 (B). UCDN refers to the WT sequence. The first row (CDN, UDN, UCN and UCD) refers to mutants with one type of endocytic determinant mutated. The second row (DN, CN, CD, UN, UD, UC) represent mutants with two types of endocytic determinants mutated. Third row (U, C, D, N) refers to mutants with three types of endocytic determinants mutated. The WT motif sequence along with the mutations introduced are shown on the right. Please refer to text for details

**Fig. 12.**
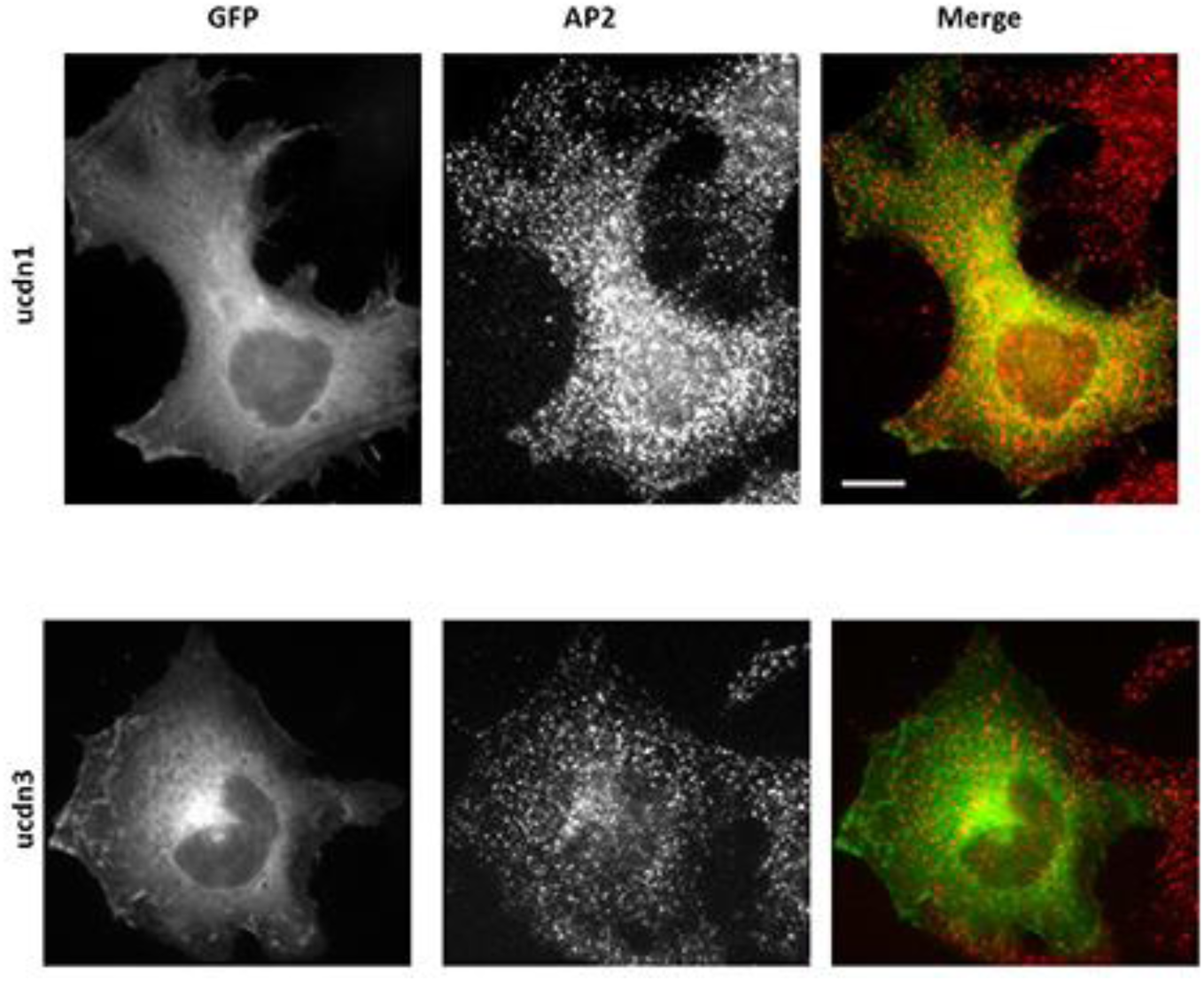
The endocytic determinants in epsin 1 and 3 are required for proper epsin localization. ucdn-1 and ucdn-3 have all 4 kinds of endocytic determinants mutated in FL epsin1 and epsin3 respectively. Scale Bar=10μm

**Fig. 13.**
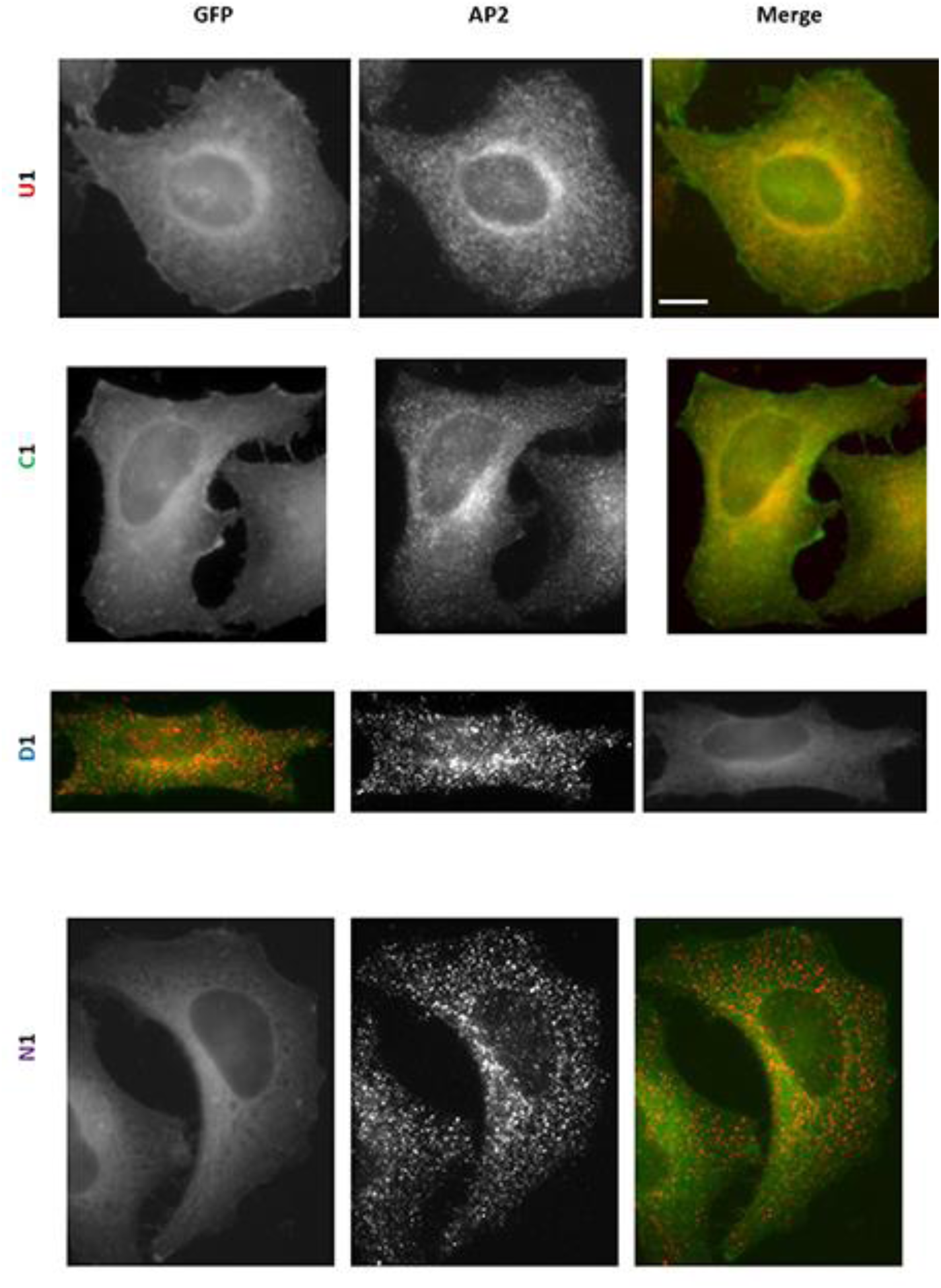
Localization of epsin1 mutants with only 1 kind of endocytic determinant intact. Representative images of the mutants stained with AP2 are shown here. Scale Bar = 10um

**Fig. 16.**
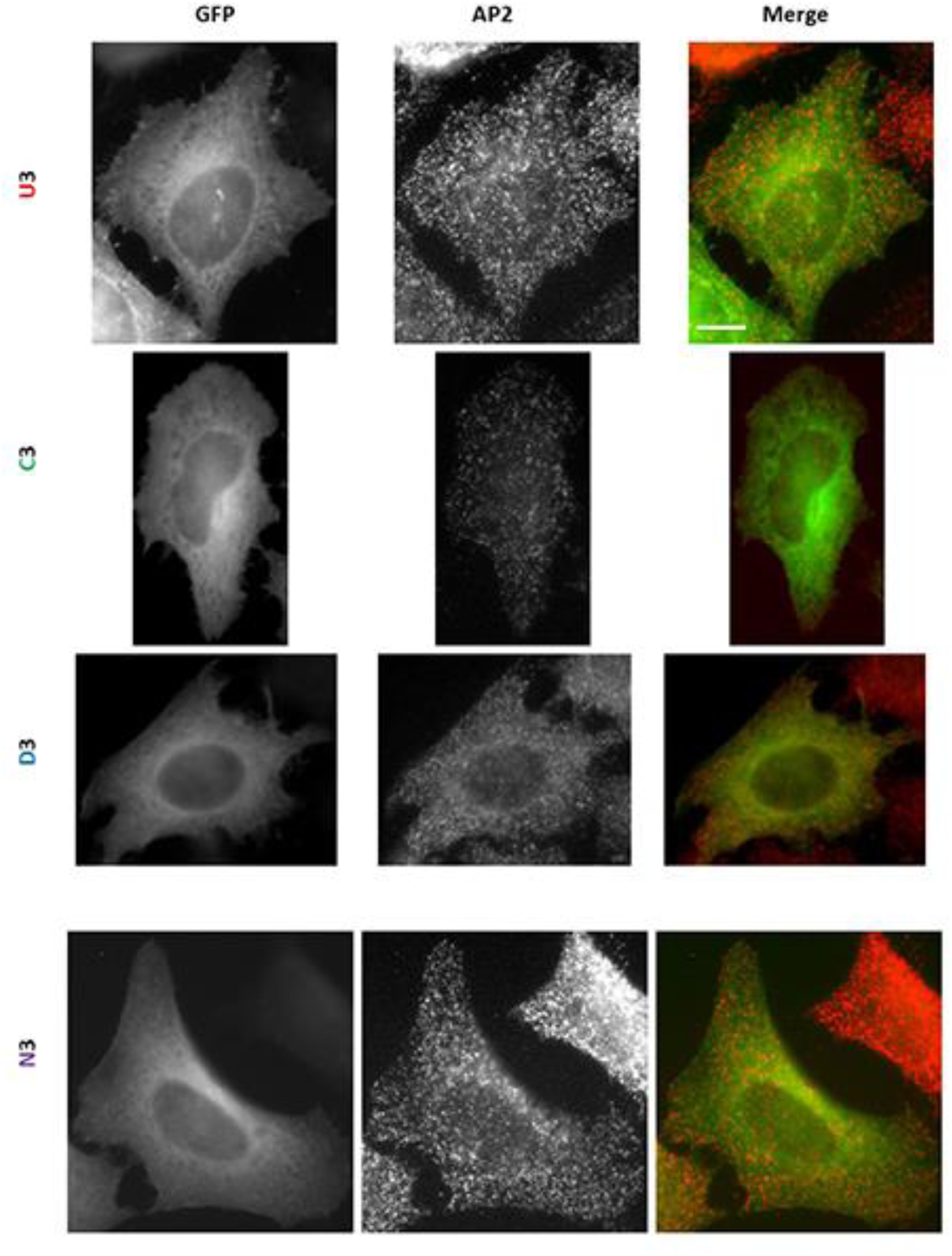
Localization of epsin3 mutants with only 1 kind of endocytic determinant intact. Representative images of the mutants stained with AP2 are shown here. Scale Bar = 10um

The next set of mutants with 2 types of endocytic determinants namely DN, CN, CD, UN, UD, UC intact gave us the most information about the hierarchy for epsin2. Recall that in Ct epsin2 mutants we observed localization dependent on the CBMs. Epsin3 Ct mutants lacking CBMs namely Ct-DN3, Ct-UD3 and Ct-UN3 could not localize at endocytic sites (Fig.17). Mutants of this category in FL epsin3 localized at endocytic sites, the most affected being UN3 (Fig.18). However, in epsin1 mutants, different localization pattern was observed. None of the 6 mutants namely, Ct-DN1, Ct-CN1, Ct-CD1, Ct-UN1, Ct-UD1, Ct-UC1 mutants could localize at endocytic sites (Fig.19). Comparing these mutants with their FL counterparts revealed 2 categories showing different localization pattern dependent on the ABMs. Mutants with ABMs and another kind of endocytic determinant mutated (CN1, UN1 or UC1) could not localize to endocytic sites, while the other 3 with intact ABMs (DN1, CD1, UD1) could localize well (Fig.20). These results suggest that while CBMs are the strongest localization determinants of epsin2 and 3, epsin1 is more dependent on ABMs for localization to endocytic sites.

**Fig. 14.**
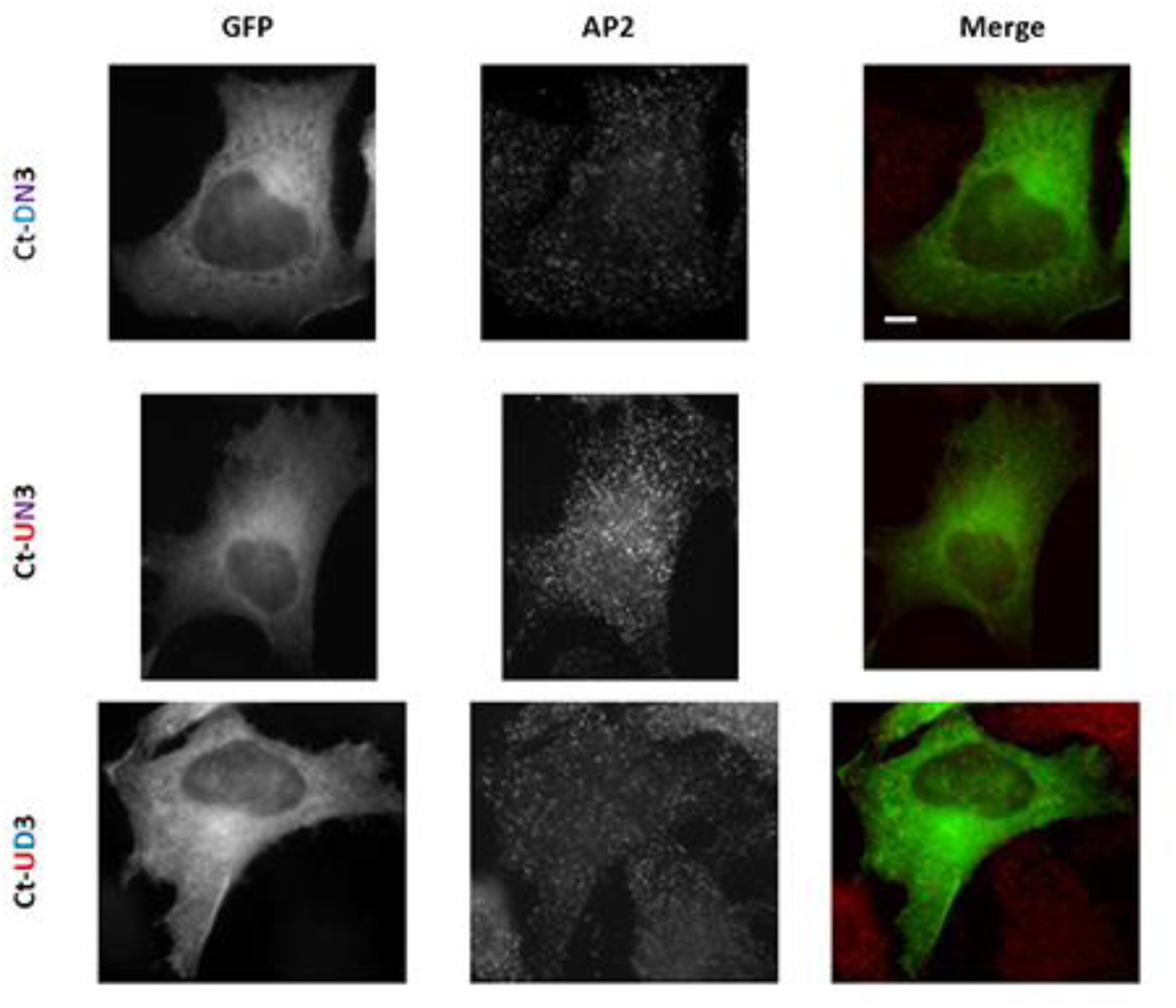
CBM is the strongest localization determinant in Epsin3. Representative images of the mutants with 2 kinds of endocytic determinants in the context of Ct-3 are shown. These mutants lack CBMs along with another kind of endocytic determinant. Scale Bar = 10um

**Fig. 15.**
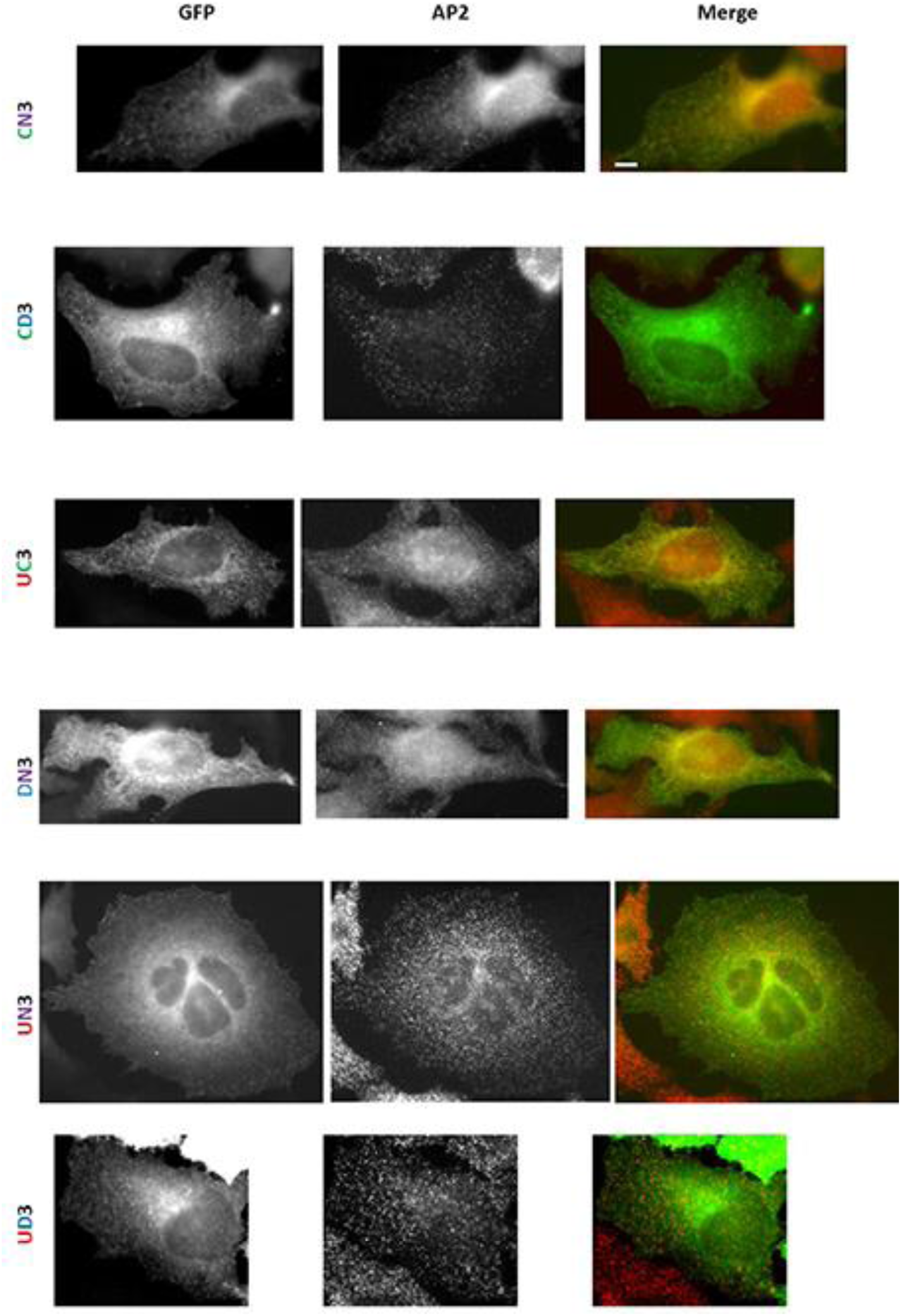
CBMs and ABMs are the strongest localization determinants in Epsin3. Representative images of the mutants with 2 kinds of endocytic determinants in the context of FL epn3 are shown. Scale Bar = 10um

**Fig. 16.**
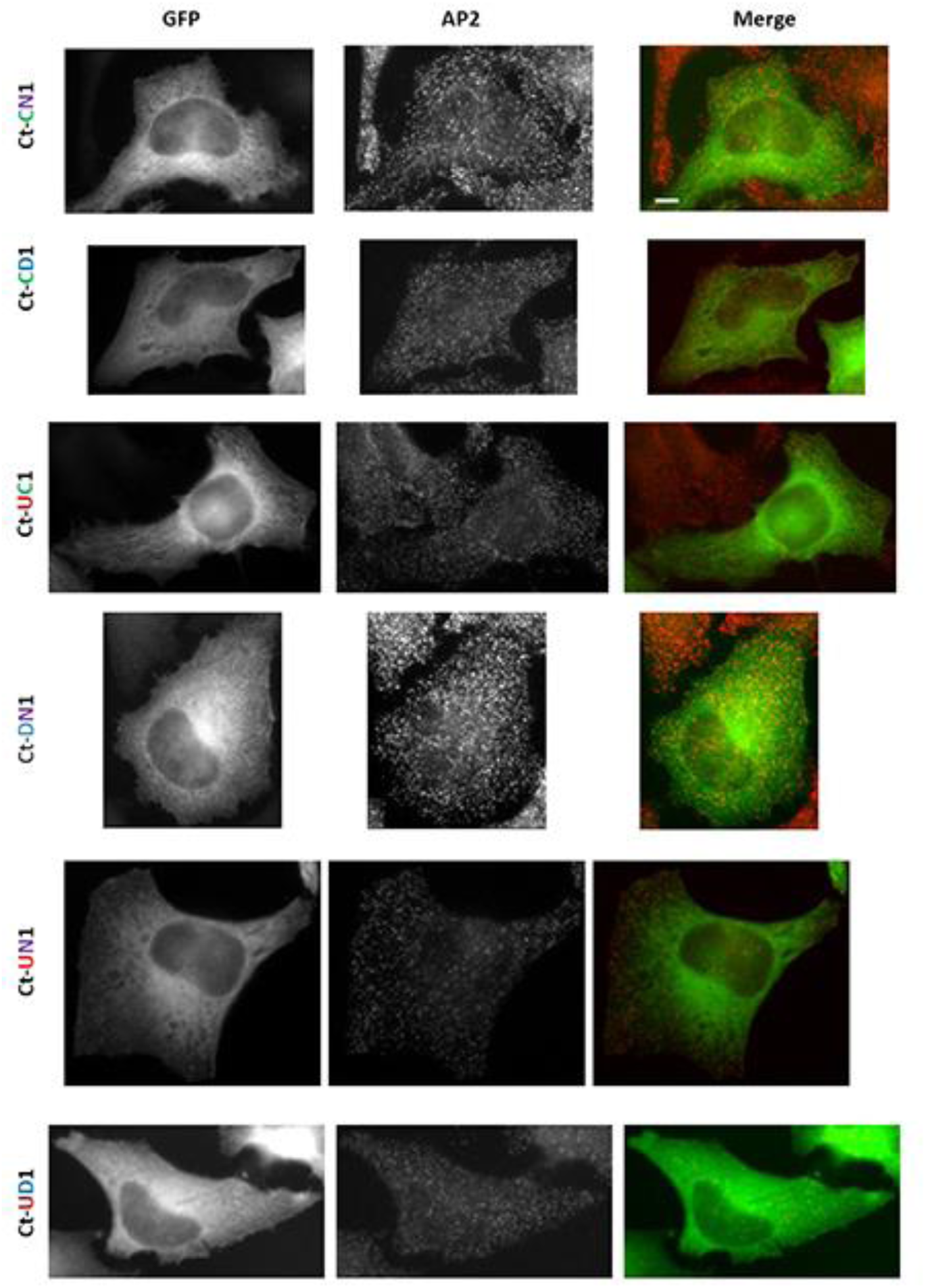
Epsin1 is the weakest paralog. Representative images of the mutants with 2 kinds of endocytic determinants in the context of Ct-1 are shown. Scale Bar = 10um

**Fig. 17.**
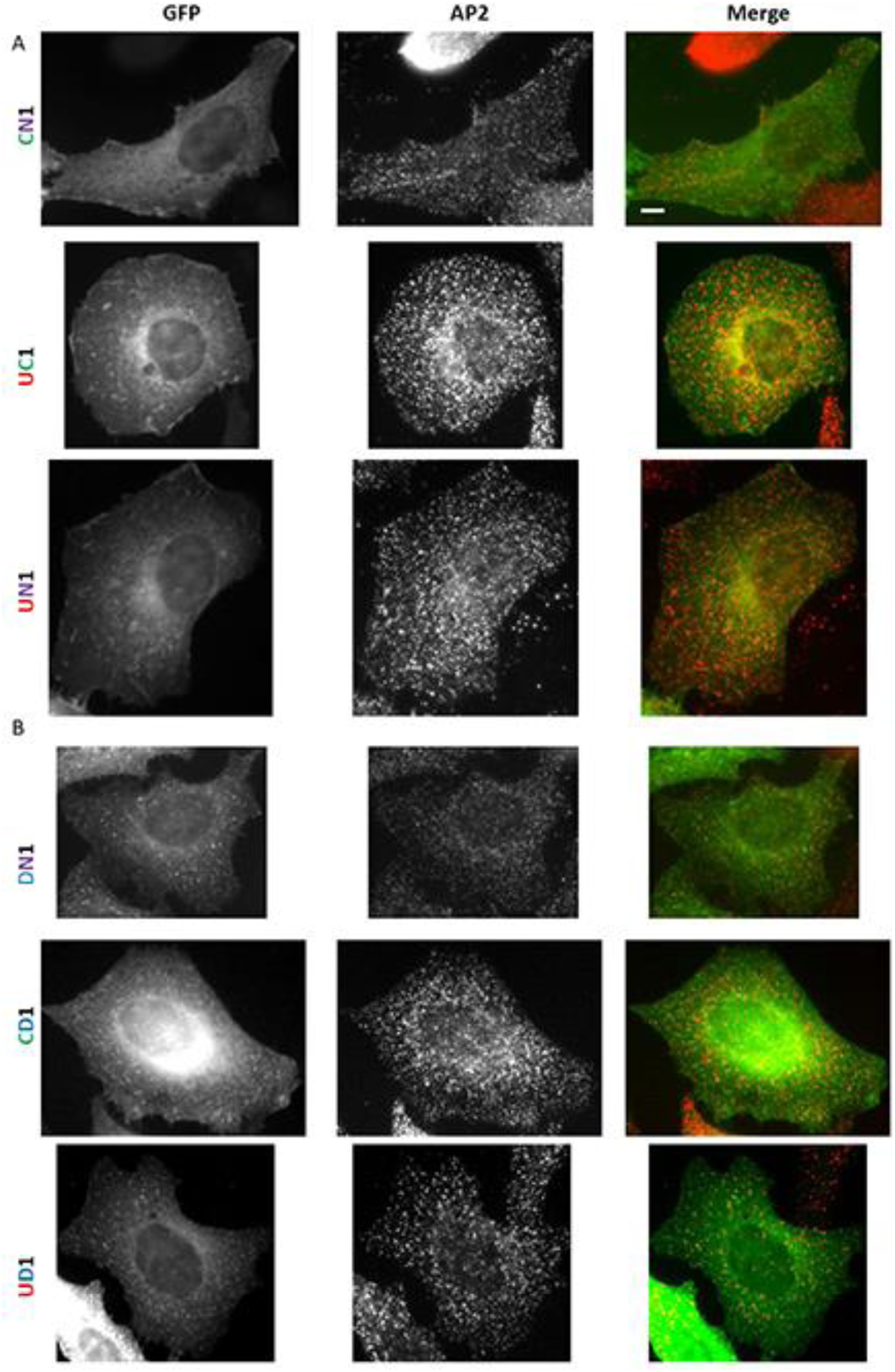
ABMs are the strongest localization determinant of epsin1. Representative images and quantification of the mutants with 2 kinds of endocytic determinants in the context of FL epn1 are shown here. A) These mutants lack ABMs along with another kind of endocytic determinant. B) These mutants have intact ABMs along with one other kind of endocytic determinant. Scale Bar = 10um

The comparison of mutants in which only 1 determinant was mutated did not show any significant difference from WT epsin2 or the Ct-2 counterparts. Epsin3 gave results very similar to epsin2, where the 4 mutants CDN3, UDN3, UCN3 and UCD3 and their C-terminal counterparts did not show any significant defect in co-localization with AP2 (Fig.21, Fig.22). However, in the case of epsin1, none of the 4 mutants Ct-CDN1, Ct-UDN1, Ct-UCN1 and Ct-UCD1 could localize well to endocytic sites (Fig.23). Nevertheless, upon comparison of their FL counterparts, only the ABMs lacking mutant UCN1 showed significant defect in co-localization with AP2 (Fig.24). While epsin3 functions very similar to epsin2, epsin1 seems to be the weakest of the paralogs, since none of the mutants could localize properly in the absence of the ENTH domain.

**Fig. 18.**
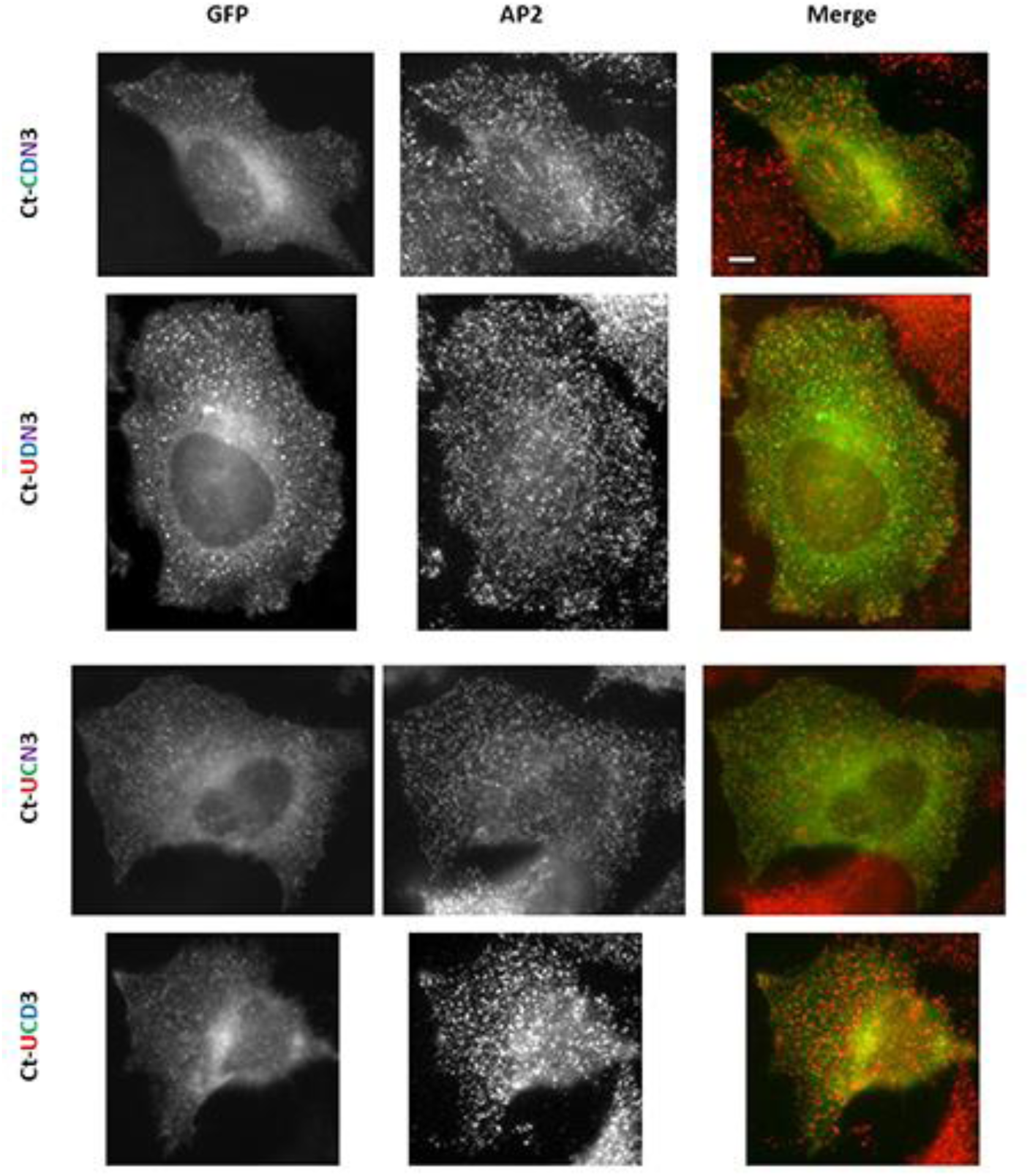
Redundancy and co-operation among the different endocytic determinants. Representative images and quantification of the mutants with 3 kinds of endocytic determinants in the context of Ct-3 are shown. Scale Bar = 10um

**Fig. 19.**
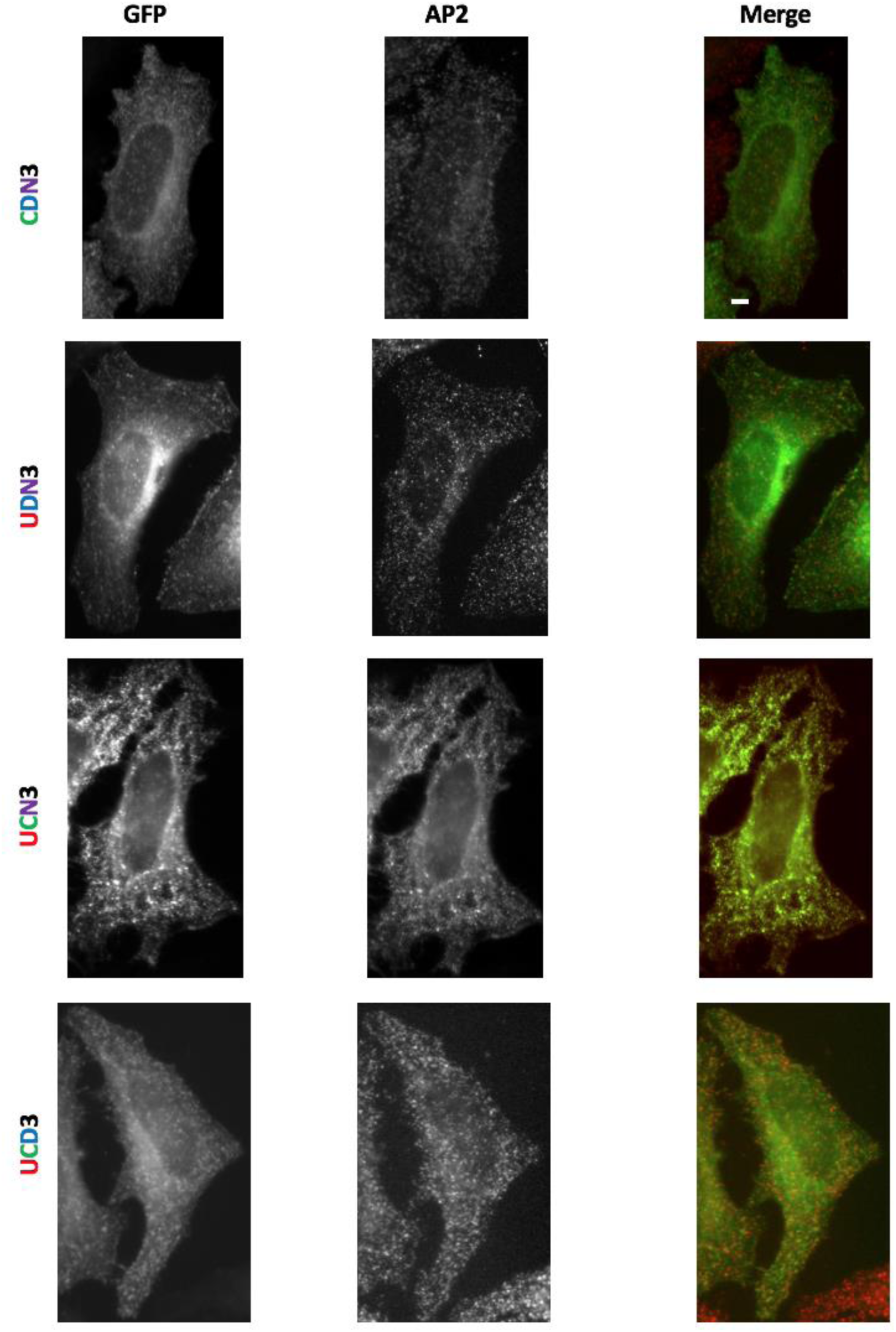
Redundancy and co-operation among the different endocytic determinants. Representative images of the mutants with 3 kinds of endocytic determinants in the context of FL epsin 3 are shown. Scale Bar = 10um

**Fig. 20.**
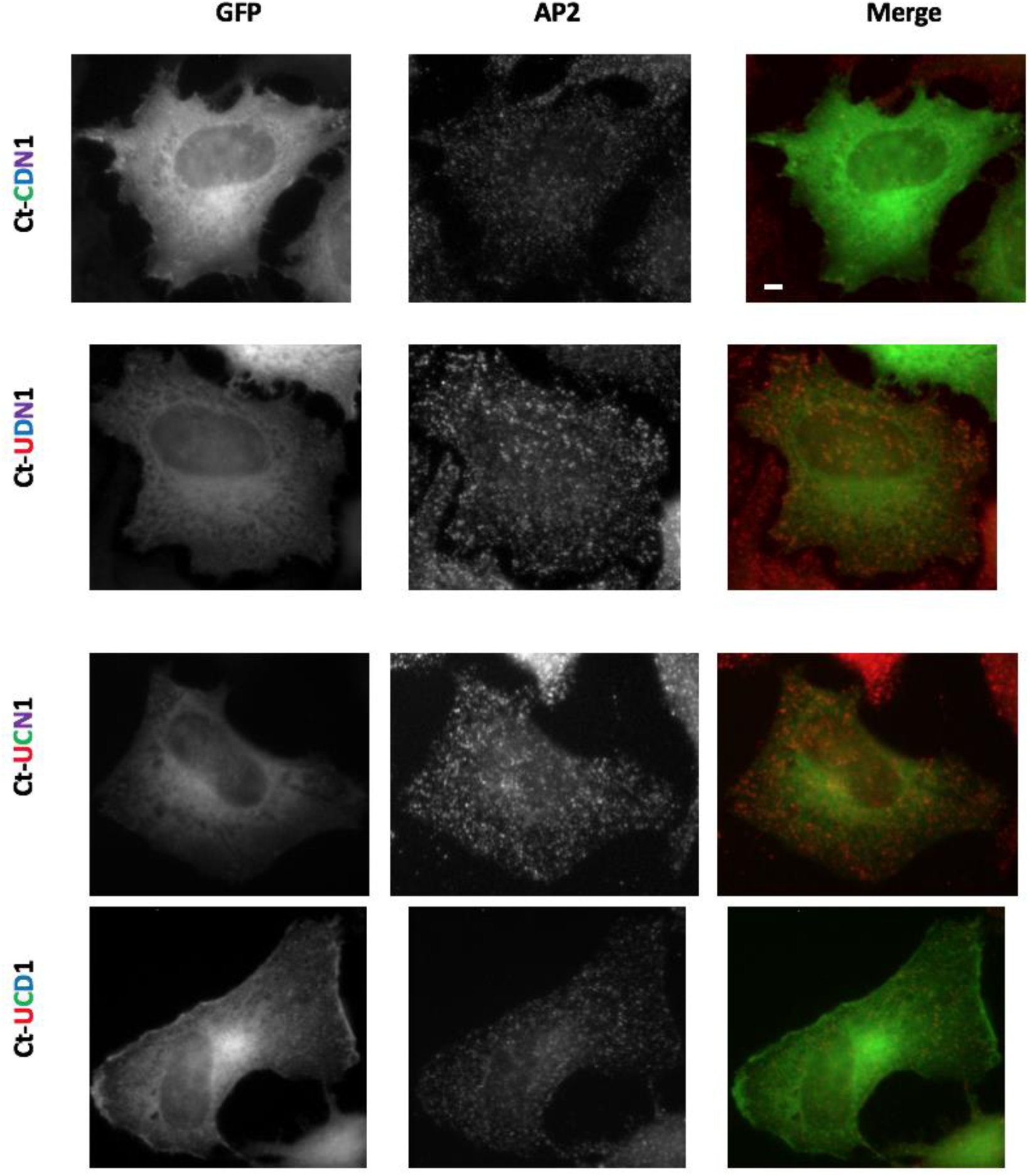
Epsin 1 is the weakest paralog. Representative images of the mutants with 3 kinds of endocytic determinants in the context of Ct-1. Scale Bar = 10um

**Fig. 21.**
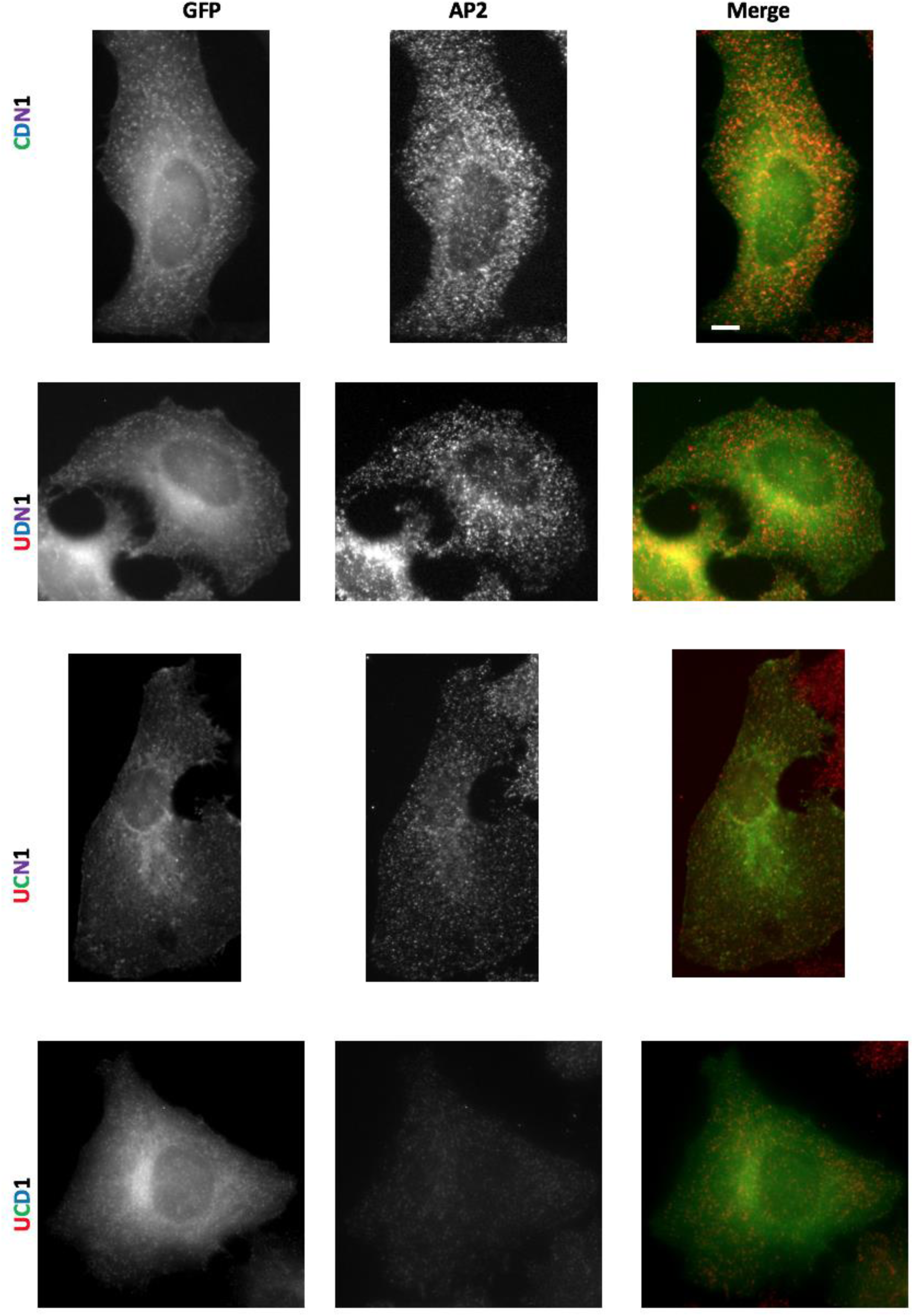
Redundancy and co-operation among the different endocytic determinants. Representative images of the mutants with 3 kinds of endocytic determinants in the context of FL epsin 1 are shown. Scale Bar = 10um

The only mutant showing substantial localization difference between epsin2 and epsin3 is UN, where UN2 could not localize to endocytic sites, while UN3 showed very low co-localization with AP2. This suggested that epsin3 probably has more determinants which could still contribute to localization other than CBMs and ABMs. So, we explored the possibility of the paralog specific region identified in epsin3 in contribution to localization.

### Role of the paralog specific sequences for epsin localization

To test the contribution of the epsin3 specific region, we deleted that region in FL-epsin3 (Epn3P), but saw no difference in co-localization with AP2, in comparison to WT epsin (Fig.25). Further, epsin3 region deleted in mutants where three kinds of endocytic determinants were present also did not show any defect in localization (Fig.25). Expanding the mutants, we also deleted the epsin3 specific region in mutants with only 2 kinds of endocytic determinants (DN3, CN3, CD3, UN3, UD3, UC3) showed a decrease in AP2 co-localization (Fig.26). The most affected mutant was UN3P, which now lacked both the strong localization determinants and the epsin3 specific region. This shows that the paralog specific region present in epsin3 makes it more competitive for localization at endocytic sites in comparison to epsin1 and epsin2. These findings imply that the strongest epsin, in terms of localization follow the order: epsin 3> 2> 1. Notably, this is also the order in which the different paralogs enhance migration.

**Fig. 22.**
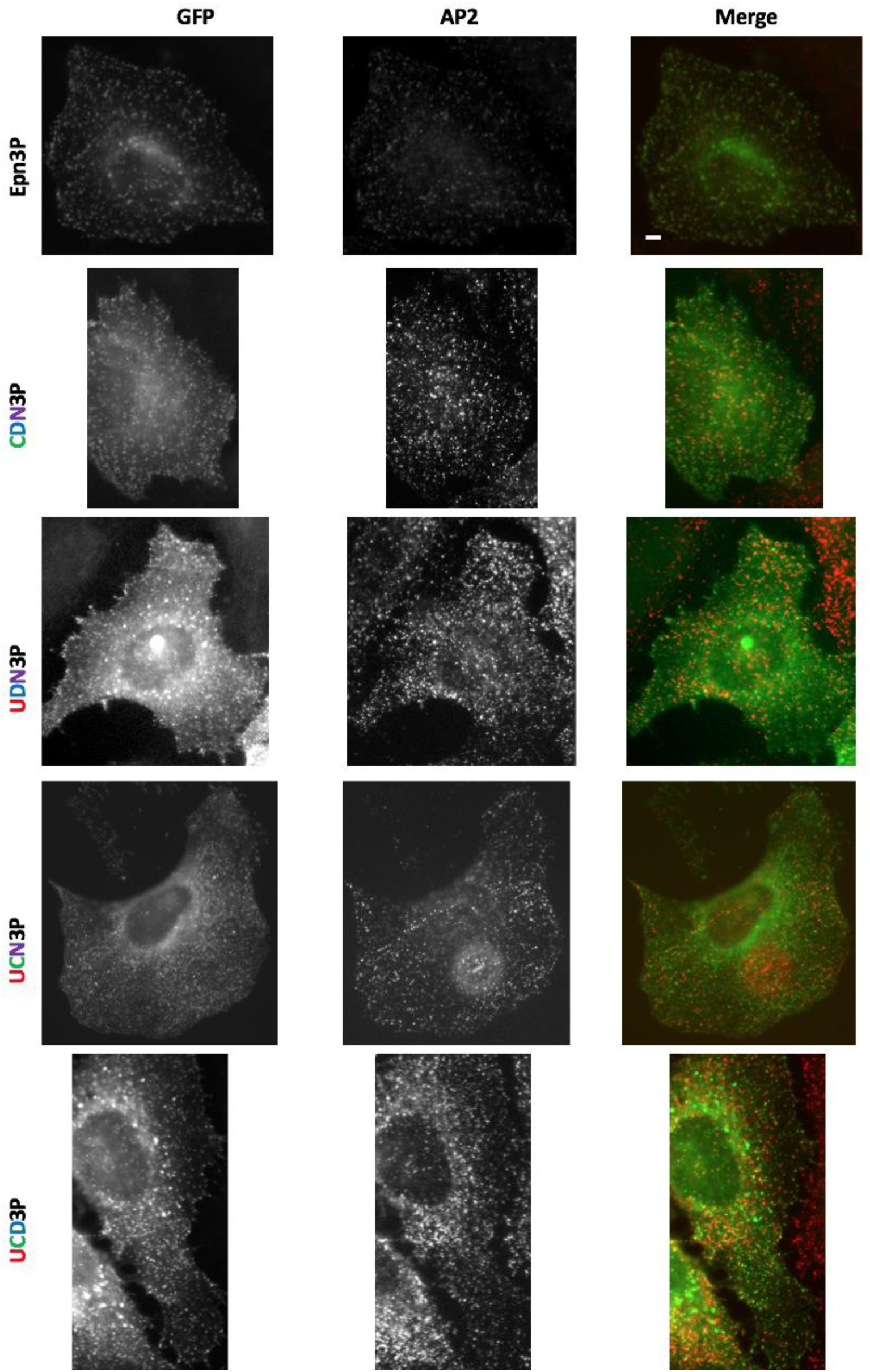
Contribution of the epsin3 specific region to localization. Representative images of the mutants with 3 kinds of endocytic determinants and the epsin3 specific region deleted in the context of FL epsin3 are shown. Scale Bar = 10um

**Fig. 23.**
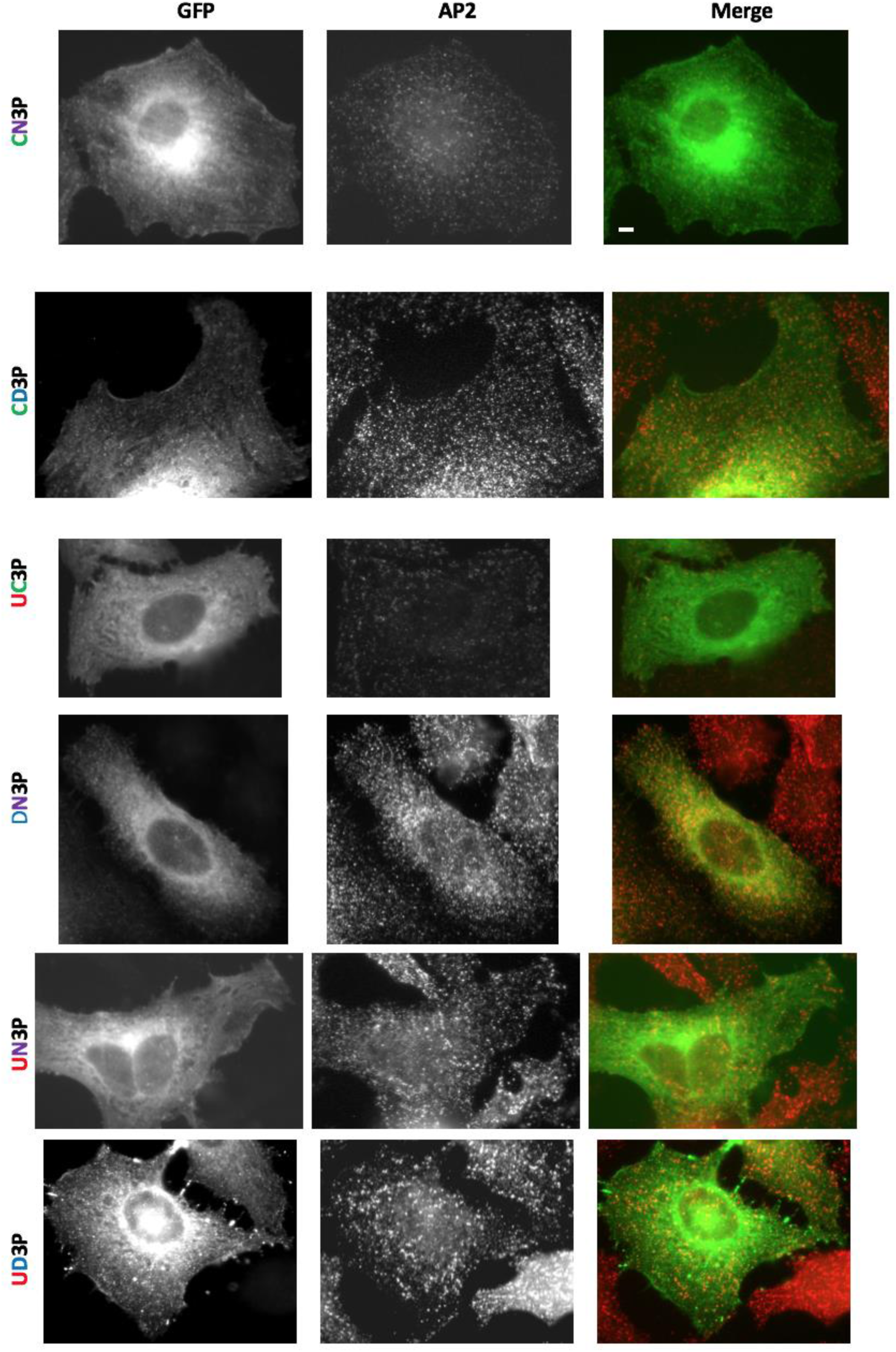
Contribution of the epsin3 specific region to localization. Representative images of the mutants with 2 kinds of endocytic determinants and the epsin3 specific region deleted in the context of FL epsin3 are shown. Scale Bar = 10um

### Role of epsin C-terminus in enhancing cell migration

Since we had established the role of the C-terminus in epsin localization, we wanted to test if a decrease in the co-localization of the epsin mutants with AP2 can be correlated with the mutant’s ability to enhance migration. We started with the FL mutants of epsin2, as it is already known that the ENTH domain is required for sustaining basal migration. So, we overexpressed the FL epsin2 mutants and assessed the ability of the cells to migrate (Fig.27). We saw a similar effect between the ability to enhance cell migration and ability to sustain proper localization in the mutants with one kind of endocytic determinant mutated and more than two kinds of endocytic determinant mutated. In the former, localization and the ability to enhance migration is retained whereas in the latter case, both are decreased. However, no similarity was observed in the mutants where 2 kinds of endocytic determinants are mutated. Consider the two mutants CD (significantly localizes at endocytic sites) and UN (completely lost localization) which have very different localization patterns, they both have lost the ability to enhance migration. We chose some representative mutants to confirm the localization versus enhancement in migration correlation in epsin3. We saw a trend very similar to epsin2. The results indicate that the reduction in the ability to enhance cell migration does not completely depend on the ability of the epsin mutants to localize at endocytic sites. Since epsin is a checkpoint protein, we rationalized that the productivity of endocytosis might be affected by mutant epsin.

**Fig. 24.**
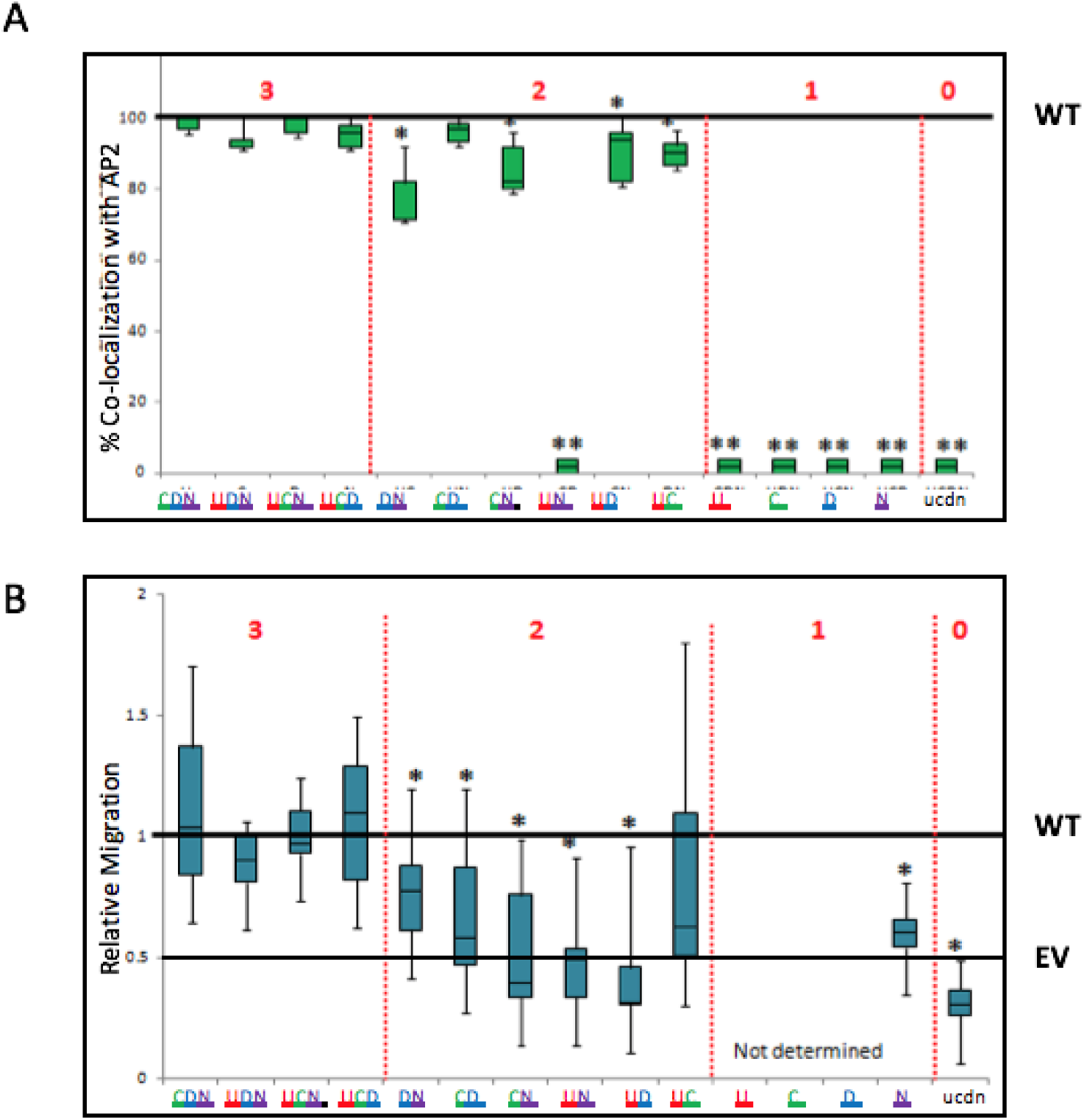
Contribution of the epsin C terminal determinants to enhancement in migration. A) This graph shows the quantification of co-localization of epsin2 FL mutants with AP2. B) This graph shows the ability of the different mutants to enhance cell migration with respect to epsin2. WT: wild-type, EV: empty vector (GFP-alone).

### Localization of epsin2 mutants in endocytic pits versus plaques

Endocytic structures can be categorized into two types, namely endocytic pits and endocytic plaques (Saffarian et al., 2009). Endocytic pits are smaller structures with shorter lifetimes of about 69sec seen in the dorsal part of the cell, while endocytic plaques are larger structures with longer lifetimes of 3 mins (approx) on the surface of the cell attached to the coverslip. Epsin has been shown to be a part of both types of endocytic structure. The function of the pits is endocytosis of specific cargoes and constant membrane turnover in the leading edge of migratory cells (Kural et al., 2015, Saffarian et al., 2009). However, the presence of plaques is cell-dependent and is linked to cell-adhesion, where a loss of plaques resulted in an increase in cell migration (Saffarian et al., 2009, Kural et al., 2015). We rationalized that if a higher proportion of the epsin (mutant) is localized to plaques in comparison to the WT it would lead to an increase in adhesion and hence cannot contribute to an enhancement in cell migration.

The previous co-localization analysis was done with images obtained with an epifluorescence scope and hence did not give us the ability to differentiate between the dorsal and ventral surfaces of the cell. We now used a spinning disk confocal microscope to measure the ability of different mutants to localize at the pits versus the plaques. Transfected cells were imaged on the dorsal versus the ventral side to quantify the density of the pits versus the plaques respectively (number of plaques or pits per unit area). As reported before by others, we observed that the dorsal and the ventral Hela cell surfaces were enriched for pits and plaques, respectively. The cells were also stained with AP2 to assess if the mutants affected the total number of endocytic sites in the cell. The results from the chosen mutants suggested that the total number of endocytic sites (no. of AP2 puncta) is not affected by epsin (or mutant) overexpression (Fig.28). However, the ability of the different mutants to localize at pits versus the plaques was affected equally, i.e. there was no shift in proportion from pit versus plaque localization. So, we turned our attention to the endocytic pits, as their activity was more important for cell migration

**Fig. 25.**
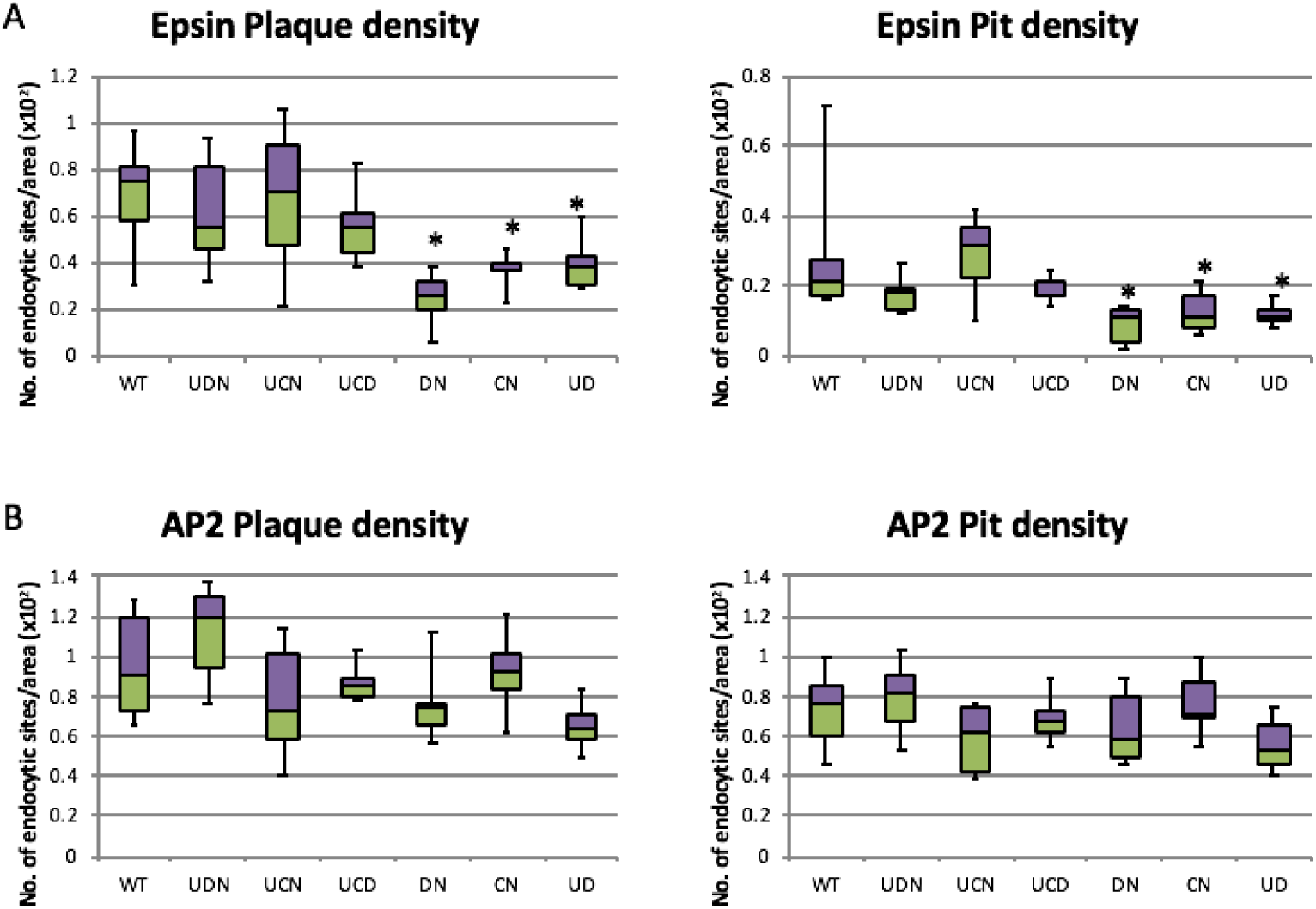
Pits versus plaque localization of epsin2 mutants. A) The graph represents the quantification of epsin 2 WT and mutant localization in pits/ plaques. B) This graph represents theAP2 density in the dorsal versus the ventral surface of the cells transfected with different epsin mutants. ** signifies p≤0.01 by Wilcoxon test in comparison to WT epn2 transfected cells. Purple and green represent first and third quartiles respectively separated by the median (second quartile).

### Epsin mutants result in more abortive pits

An initiated endocytic pit can have 3 different fates leading to an abortive, productive or a persistent pit (Loerke et al., 2009). The three dynamically distinct pits have different lifetimes of 5 to 30sec for abortive pits, approx 90 secs for the productive pits and greater than 90sec for persistent pits. Several accessory proteins were evaluated for their ability to shift the balance between the proportions of abortive versus productive sites (Mettlen et al., 2009b). Epsin, among others, emerged as a member which imposes a ‘checkpoint’ in determining the productivity of the endocytic sites. Epsin has a unique ability to interact with multiple components of the endocytic machinery including clathrin, AP2, ubiquitinated cargo, several accessory proteins and actin (Sen et al., 2012). We then tested transfected cells with chosen epsin mutants and studied the dynamics of the endocytic sites that they are localized to. The results clearly show that epsin mutants alter the ratio of productive to non-productive (abortive and persistent) sites (Fig.29). The decrease in productive and increase in non-productive events was exaggerated in epsin mutants as compared to the WT. Moreover, the differences were higher in the mutants with only 2 endocytic determinants versus the mutants with 3 endocytic determinants.

**Fig. 26.**
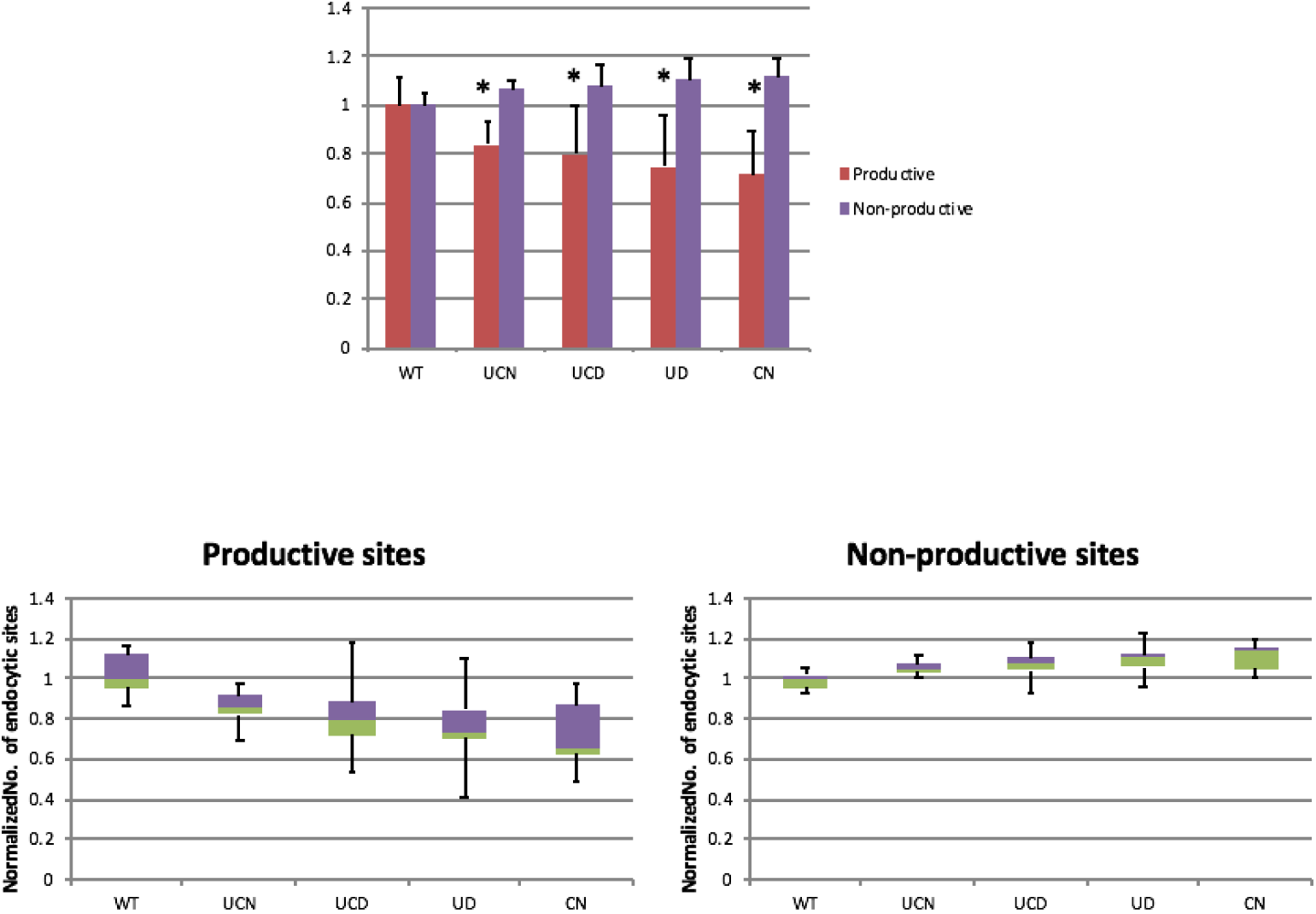
Productivity of endocytic sites affected by epsin mutants. The graph shows the number of productive and non-productive endocytic sites as measured normalized to WT epsin. * signifies p<0.05 by t-test.

## CONCLUSIONS AND DISCUSSION

Endocytosis is an intricately orchestrated, vital cellular process. Endocytosis is required for nutrient uptake, receptor downregulation, neurotransmission and polarity dependent processes such as cell migration and invasion (McMahon and Boucrot, 2011; Saheki and De Camilli, 2012; Cosker and Segal, 2014; Caswell and Normal, 2008). Endocytosis consequently contributes to the regulation of several signaling events in space and time. Signaling can also trigger endocytosis downstream resulting in the two intertwining networks: signaling and endocytosis (Di-Fiore and von Zastow, 2014; Sorkin and Von Zastrow, 2009). This work studied the synergism of Epsin ENTH domain with its signaling function and and the C-terminus endocytic functions in enhancing cell migration.

Epsin ENTH domain binds to PI(4,5)P_2_ enriched in the plasma membrane and the C-terminus binds to different components of the endocytic machinery. There has been a long-term ambiguity in the localization determinant of Epsin. So, to test this, we used the ENTH-GFP and Ct-GFP (del ENTH) and studied its ability to localize at endocytic sites (marked by AP2). We showed that while the ENTH domain is required for defining the specificity of epsin localization, the major determinant of localization are the endocytic determinants within the C-terminus. Specifically, in the absence of the ENTH domain, while localization at endocytic sites was not significantly affected, an enrichment of epsin was observed in the TGN, which was lost upon mutation of the CBMs. This suggested that epsin was binding to clathrin at the TGN in the absence of the lipid binding domain (ENTH), which targets it to the cell membrane. This suggested that the C-terminus of Epsin was the major determinant of localization with the ENTH domain playing a supporting role. We used a systematic combinatorial approach to create a comprehensive mutation collection of the 4 kinds of endocytic determinants to determine the contribution of the individual endocytic determinants to epsin localization. Epsin 2 was the first paralog to be studied because of the commonalities it shared with the other paralogs. The results are summarized below.

Endocytic determinants in the C-terminus which comprise less than 10% of the sequence are required for proper epsin2 localization.
At least two different kinds of endocytic determinants are needed for proper epsin2 localization. This reflects co-operativity within the different endocytic determinants.
CBMs are the strongest epsin2 localization determinant. Specifically, the distal CBM2 which followed the classical CBM consensus (LФpФp) was more important for localization.
Endocytic determinants in decreasing order of importance for epsin2 localization: CBM > DPW > NPF ≥ UIM
ENTH domain reinforces the effect of the weak determinants on epsin2 localization (obtained by comparing the mutants UD2 and DN2 in FL and Ct context)

Three mammalian epsin paralogs (epsin 1,2 and 3) have been identified. Functional redundancy of epsin1 and 2 has been reported previously (Chen et al., 2009, Tessneer et al., 2013, Messa et al., 2014). Epsin3 is found specifically upregulated in invasive cancers (Coon et al., 2011). Our lab previously reported that the epsins have differential abilities to enhance cell migration and invasion, suggesting that there are some inherent differences between the 3 paralogs (Coon et al., 2010). However, all 3 paralogs localize to endocytic site similarly. So, we wondered if the difference in localization determinants could explain their differential ability to enhance migration. So we created key mutants in the other paralogs and performed localization analysis.

Indeed, they exhibited some difference in the hierarchy of the endocytic determinants. While epsin2 and 3 followed a similar hierarchy (CBMs>ABMs>UIMs≥NPFs), epsin1 (ABMs>CBMs≥UIMs≥NPFs) was different. Furthermore, epsin2 and 3 mutants, but not epsin 1 could localize at endocytic sites with just 3 types of endocytic determinants in the absence of the ENTH domain. This suggested that epsin2 and 3 are more competent for recruitment to endocytic sites than epsin 1. Comparing UN2 and UN3 showed that epsin 3 mutants could still barely localize to endocytic sites, while epsin 2 could not. UN3P obtained by deleting the epsin 3 specific region completely lost the ability to localize at endocytic sites. Collectively, these finding suggested that there were some inherent differences between the three epsin paralogs based on the differences found in their C-terminus. More specifically, epsin1 was the weakest paralog among the 3. Based on the determinants present in the 3 different paralogs contributing to localization, Epsin3>2>1, which is incidentally the order in which the Epsins enhance cell migration.

After establishment of the localization determinants, we wanted to determine if the mutants’ efficiency in endocytic site localization paralleled its ability to enhance cell migration. We already established that the ENTH domain is essential for cell migration, so we used only FL mutants for the migration assays. We were surprised to see that even though some mutants could localize to endocytic sites with 70-80% efficiency, they were unable to enhance cell migration. The mutants which completely lost the ability to localize at endocytic sites could not enhance cell migration. However, even though epsin2 mutants with 2 types of endocytic determinants could localize well, they could not enhance cell migration, but fell to the level of the EV. This finding suggested that merely just the presence of strong localization determinants was not sufficient to enhance migration. So, we monitored the productivity of the endocytic sites formed by the mutants by performing live cell imaging. Previous studies have shown that endocytic sites are initiated by random sampling of the membrane by clathrin and AP2 (Ehrlich et al., 2004). Recruitment of epsin to endocytic sites has been shown to shadow clathrin and AP2 at endocytic sites (Saffarian et al., 2009). Epsin has also been shown to function as a sensor in determining the maturation of the endocytic pit (Loerke et al., 2009). In fact we found that when compared to epsin2 WT, mutants showed an increase in abortive sites and a decrease in productive sites. In fact, the productivity was progressively reduced in mutants with lesser endocytic determinants. These results clearly point out to the endocytic function of epsins contribution to enhancing cell migration.

Epsin is a unique adaptor protein which possesses the ability to bind to multiple components of the endocytic machinery including actin, clathrin, AP2, accessory proteins and ubiquitinated cargo. It has been shown to function as a specific adaptor protein for ubiquitinated cargoes VEGFR2 and EGFR which requires the integrity of its UIMs (Sen et al., 2012). However, our analysis pointed to the importance of the CBMs and DPWs for localization. This outcome was unexpected for an endocytic adaptor protein epsin which is defined by its ability to bind and internalize ubiquitinated cargo. Furthermore, just the presence of the strong endocytic determinants was not sufficient to enhance cell migration. Based on our results, we propose that Epsin goes to already initiated endocytic sites, incorporates ubiquitinated cargo and increases the size of the pit by recruiting more members (Fig.30). Eventually, only if the pit reaches a critical mass, it moves on to the maturation phase, else it’s aborted. So epsin requires all the 4 different kinds of endocytic determinants to help in maturation of endocytic pits, rather than initiation.

**Fig. 30.**
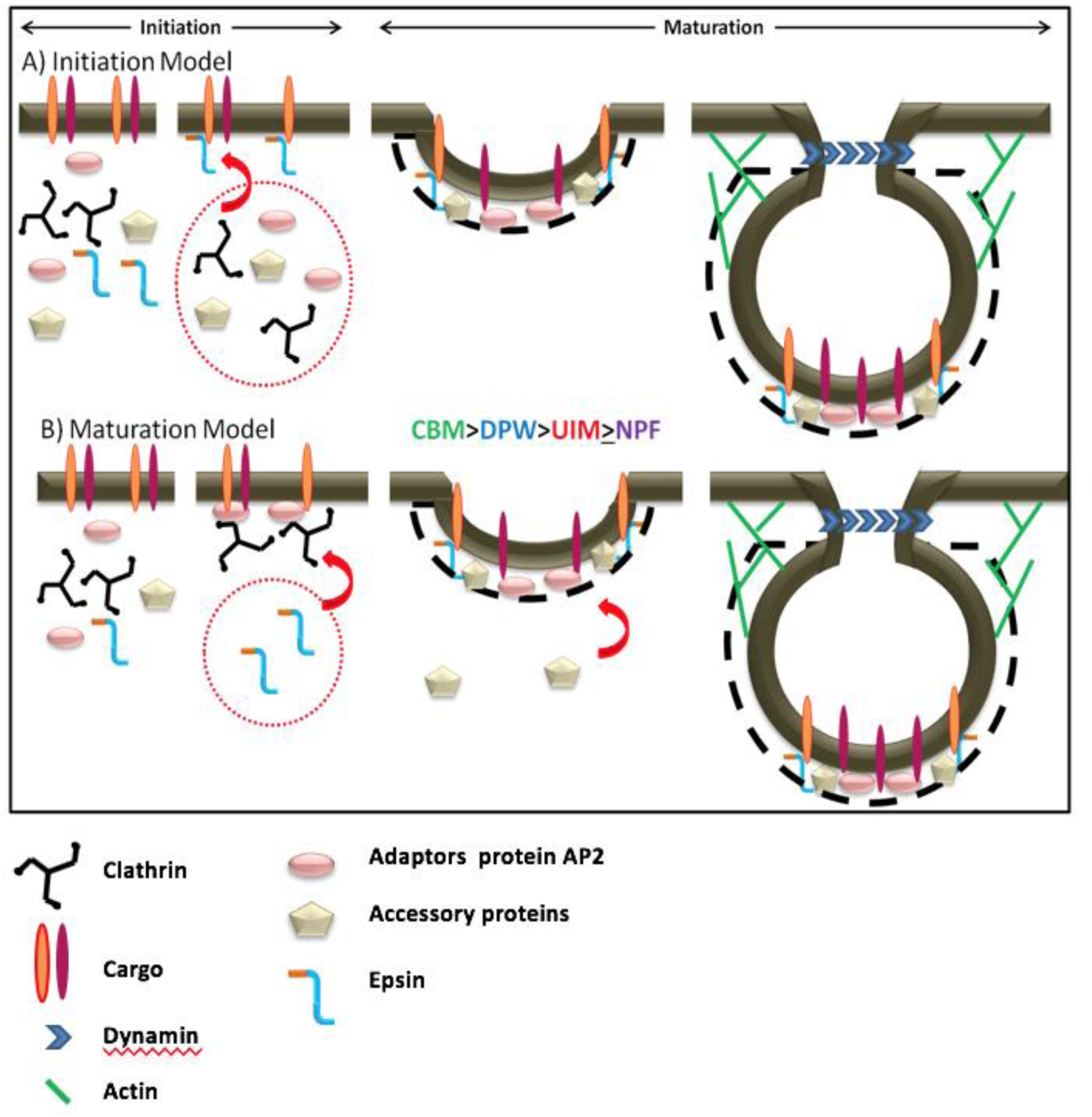
Model for the recruitment of epsin to endocytic sites. Our result support a maturation model for epsin as opposed to the initiation model. According to the maturation model, epsin recruits to an existing endocytic site and helps in the maturation process. According to the initiation model, epsin initiated an endocytic site around an ubiquitinated cargo.

## MATERIALS AND METHODS

### Cells and Culture conditions

Hela cells were cultured in DMEM, Streptomycin/Penicillin, 2mM L-Glutamine and 10% fetal bovine serum (FBS) at 37 °C in a 5% CO_2_ incubator.

### Quantification of co-localization

We developed a method of quantifying co-localization of Epsin (GFP-tagged construct) with AP2 (marker of endocytic sites). Random images totaling approximately 200 punctate structures in least 5 cells were analyzed for each mutant and were repeated twice. Single-stained cells were used to estimate the amount of signal detected in channel “A” that corresponds to non-filtered fluorescence from channel “B” (crosstalk). Channel “A” crosstalk vs. fluorescence intensity in channel “B” curves were constructed and subjected to regression. These functions were used to set up thresholds (mean crosstalk+1xSD) above which co-localization signals in dual stained cells will be considered to be statistically significant. Also only the fluorescence intensity values above the cytoplasmic fluorescence intensity in Channel “A” were considered as puncta. Finally percentage co-localization was calculated using the formula below,

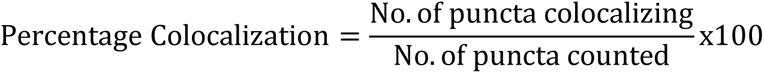

To measure nuclear and TGN localization, we stained the cells with DAPI or TGN marker (TGN46) respectively along with the mutant (GFP-tagged) to be studied. Tracing the nucleus in the DAPI channel or the TGN in its respective channel and measuring the intensity in the GFP channel gave the nuclear and TGN intensities in the green channel. Tracing the perimeter of the cell and measuring the intensity in the GFP channel gave the total cell intensity. Tracing a region of the cell devoid of puncta gave us the cytoplasmic intensity of the cell which when integrated over the area gave the total cytoplasmic intensity. Finally we had 4 components for each mutant: percentage of co-localization with AP2, percentage of localization in the nucleus, percentage of localization in the TGN and percentage of localization in the cytoplasm. These values were represented as a bar graph for every mutant. Puncta and region tracing was done manually using the Image J software.

### Migration Assay

In this assay, 10^4^ cells were trypsinized and trypsin quenched with complete media. Cells were then resuspended in 500µl media lacking FBS and allowed to recover in suspension for 1 h. Then cells were applied on upper chamber of the 0.33 cm^2^, 8 µm pore transwell inserts (Corning Inc) coated with 100 µg/ml BSA and 10 µg/ml fibronectin on the upper and bottom side of the insert membrane, respectively. The migration chambers were then placed in wells containing media with 10% FBS and allowed to migrate for 3 h and fixed in 3% formaldehyde for 10 min. Cells on fixed membranes were then stained with DAPI and visualized by epifluorescence microscopy. The number of transfected cells per membrane was counted in order to obtain cell inputs, and then the upper side of the membrane was wiped with a cotton swab and rinsed with PBS. Cells on the bottom side of the membrane were then counted and scored as migrants if their nuclei passed through the membrane pores. Percentage migration was determined as the ratio of migrated cells to the total number of transfected cells. Migration results were normalized based on WT percentage migration and expressed as mean ± SD of at least triplicate membranes.

### Dynamics of endocytic cells

Cells were transfected with GFP-tagged Epsin WT or epsin mutants and seeded in live cell imaging chambers 12 hours before the experiment. After attaching, they were starved for 4 hours. Just before imaging, they were stimulated with serum containing media and sealed with parafilm. Imaging was done using a spinning disc confocal microscopy for 30 mins. Each cell was imaged for 3 mins on the dorsal surface at an interval of 1 sec. 6 movies per sample were obtained. The movies were analyzed using the Image J plugin Speckle Tracker J (Smith et al., 2011). Only pits that were initiated after frame 1 and which disappeared before the last frame were considered for the analysis. 500-1000 spots per mutant were analyzed.

## CONSTRUCTS USED IN THIS STUDY

CONSTRUCTS USED IN THIS STUDY:
| Plasmid | Description | Source |
| --- | --- | --- |
| GFP | pEGFP-C1 | Clontech |
| Epn1-GFP | *Rattus norvegicus* Epsin1 in pEGFP-N1 | H.McMahon (MRC Cambridge) |
| Epn2-GFP | *Homo sapiens* Epsin2a in pEGFP-C1 | P. De Camilli (Yale) |
| Epn2-mCherry | *Homo sapiens* Epsin2 in mCherry-C1 | Taylor et al, 2011 |
| Epn3-GFP | *Homo sapiens* Epsin3 in pEGFP-C1 | B.Pilcher (Bioform Medical) |
| ENTH1-GFP | *Rattus norvegicus* Epsin1<sub>1-168</sub> pEGFP | Coon et al., 2010 |
| ENTH2-GFP | *Homo sapiens* Epsin2<sub>1-160</sub> pEGFP | This study |
| Ct1-GFP | *Rattus norvegicus* Epsin1<sub>166-550</sub> pEGFP | Coon et al., 2010 |
| Ct2-GFP | *Homo sapiens* Epsin2<sub>160-640</sub> pEGFP | This study |
| Ct3-GFP | *Homo sapiens* Epsin3<sub>176-633</sub> pEGFP | This study |
| CDN1-GFP | *Rattus norvegicus* Epsin1 UIM1&2 mutated<br>in pEGFP | This study |
| UDN1-GFP | *Rattus norvegicus* Epsin1 CBM1 &2 mutated<br>in pEGFP | This study |
| UCN1-GFP | *Rattus norvegicus* Epsin1DPW1 to 8 mutated<br>in pEGFP | This study |
| UCD1-GFP | *Rattus norvegicus* Epsin1 NPF1 to 3 mutated<br>in pEGFP | This study |
| DN1-GFP | *Rattus norvegicus* Epsin1UIMs & CBMs<br>mutated in pEGFP | This study |
| CN1-GFP | *Rattus norvegicus* Epsin1UIMs & DPWs<br>mutated in pEGFP | This study |
| CD1-GFP | *Rattus norvegicus* Epsin1 UIMs & NPFs<br>mutated in pEGFP | This study |
| UN1-GFP | *Rattus norvegicus* Epsin1 CBMs & DPWs<br>mutated in pEGFP | This study |
| UD1-GFP | *Rattus norvegicus* Epsin1 CBMs & NPFs<br>mutated in pEGFP | This study |
| UC1-GFP | *Rattus norvegicus* Epsin1 DPWs and NPFs<br>mutated in pEGFP | This study |
| U1-GFP | *Rattus norvegicus* Epsin1 CBMs, DPWs &<br>NPFs mutated in pEGFP | This study |
| C1-GFP | *Rattus norvegicus* Epsin1 UIMs, DPWs & NPFs mutated in pEGFP | This study |
| D1-GFP | *Rattus norvegicus* Epsin1 UIMs, CBMs & NPFs mutated in pEGFP | This study |
| N1-GFP | *Rattus norvegicus* Epsin1 UIMs, CBMs & DPWs mutated in pEGFP | This study |
| ucdn1-GFP | *Rattus norvegicus* Epsin1 UIMs, CBMs, DPWs and NPFs mutated in pEGFP | This study |
| Ct-CDN1-GFP | *Rattus norvegicus* Epsin1<sup>166-550</sup> UIM1&2 mutated in pEGFP | This study |
| Ct-UDN1-GFP | *Rattus norvegicus* Epsin1<sup>166-550</sup> CBM1 &2 mutated in pEGFP | This study |
| Ct-UCN1-GFP | *Rattus norvegicus* DPW1 to 8 mutated in pEGFP | This study |
| Ct-UCD1-GFP | *Rattus norvegicus* Epsin1<sup>166-550</sup> NPF1 to 3 mutated in pEGFP | This study |
| Ct-DN1-GFP | *Rattus norvegicus* Epsin1<sup>166-550</sup> UIMs & CBMs mutated in pEGFP | This study |
| Ct-CN1-GFP | *Rattus norvegicus* Epsin1<sup>166-550</sup> UIMs & DPWs mutated in pEGFP | This study |
| Ct-CD1-GFP | *Rattus norvegicus* Epsin1<sup>166-550</sup> UIMs & NPFs mutated in pEGFP | This study |
| Ct-UN1-GFP | *Rattus norvegicus* Epsin1<sub>166-550</sub> CBMs & DPWs mutated in pEGFP | This study |
| Ct-UD1-GFP | *Rattus norvegicus* Epsin1<sub>166-550</sub> CBMs & NPFs mutated in pEGFP | This study |
| Ct-UC1-GFP | *Rattus norvegicus* Epsin1<sub>166-550</sub> DPWs and NPFs mutated in pEGFP | This study |
| Ct-U1-GFP | *Rattus norvegicus* Epsin1<sub>166-550</sub> CBMs, DPWs & NPFs mutated in pEGFP | This study |
| Ct-C1-GFP | *Rattus norvegicus* Epsin1<sub>166-550</sub> UIMs, DPWs & NPFs mutated in pEGFP | This study |
| Ct-D1-GFP | *Rattus norvegicus* Epsin1<sub>166-550</sub> UIMs, CBMs & NPFs mutated in pEGFP | This study |
| Ct-N1-GFP | *Rattus norvegicus* Epsin1<sub>166-550</sub> UIMs, CBMs & DPWs mutated in pEGFP | This study |
| Ct-ucdn1-GFP | *Rattus norvegicus* Epsin1<sub>166-550</sub> UIMs, CBMs, DPWs and NPFs mutated in pEGFP | This study |
| CDN2-GFP | *Homo sapiens* Epsin2 UIM1&2 mutated in pEGFP | This study |
| UDN2-GFP | *Homo sapiens* Epsin2 CBM1 &2 mutated in pEGFP | This study |
| UCN2-GFP | *Homo sapiens* Epsin2 DPW1 to 7 mutated in pEGFP | This study |
| UCD2-GFP | *Homo sapiens* Epsin2 NPF1 to 3 mutated in pEGFP | This study |
| DN2-GFP | *Homo sapiens* Epsin2 UIMs & CBMs mutated in pEGFP | This study |
| CN2-GFP | *Homo sapiens* Epsin2 UIMs & DPWs mutated in pEGFP | This study |
| CD2-GFP | *Homo sapiens* Epsin2 UIMs & NPFs mutated in pEGFP | This study |
| UN2-GFP | *Homo sapiens* Epsin2 CBMs & DPWs mutated in pEGFP | This study |
| UD2-GFP | *Homo sapiens* Epsin2 CBMs & NPFs mutated in pEGFP | This study |
| UC2-GFP | *Homo sapiens* Epsin2 DPWs and NPFs mutated in pEGFP | This study |
| U2-GFP | *Homo sapiens* Epsin2 CBMs, DPWs & NPFs mutated in pEGFP | This study |
| C2-GFP | *Homo sapiens* Epsin2 UIMs, DPWs & NPFs mutated in pEGFP | This study |
| D2-GFP | *Homo sapiens* Epsin2 UIMs, CBMs & NPFs mutated in pEGFP | This study |
| N2-GFP | *Homo sapiens* Epsin2 UIMs, CBMs & DPWs mutated in pEGFP | This study |
| ucdn2-GFP | *Homo sapiens* Epsin2 UIMs, CBMs, DPWs and NPFs mutated in pEGFP | This study |
| Ct-CDN2-GFP | *Homo sapiens* Epsin2<sub>160-640</sub> UIM1&2 mutated in pEGFP | This study |
| Ct-UDN2-GFP | *Homo sapiens* Epsin2<sub>160-640</sub> CBM1 &2 mutated in pEGFP | This study |
| Ct-UCN2-GFP | *Homo sapiens* Epsin2<sub>160-640</sub> DPW1 to 7 mutated in pEGFP | This study |
| Ct-UCD2-GFP | *Homo sapiens* Epsin2<sub>160-640</sub> NPF1 to 3 mutated in pEGFP | This study |
| Ct-DN2-GFP | *Homo sapiens* Epsin2<sub>160-640</sub> UIMs & CBMs mutated in pEGFP | This study |
| Ct-CN2-GFP | *Homo sapiens* Epsin2<sub>160-640</sub> UIMs & DPWs mutated in pEGFP | This study |
| Ct-CD2-GFP | *Homo sapiens* Epsin2<sub>160-640</sub> UIMs & NPFs mutated in pEGFP | This study |
| Ct-UN2-GFP | *Homo sapiens* Epsin2<sub>160-640</sub> CBMs & DPWs mutated in pEGFP | This study |
| Ct-UD2-GFP | *Homo sapiens* Epsin2<sub>160-640</sub> CBMs & NPFs mutated in pEGFP | This study |
| Ct-UC2-GFP | *Homo sapiens* Epsin2<sub>160-640</sub> DPWs and NPFs mutated in pEGFP | This study |
| Ct-U2-GFP | *Homo sapiens* Epsin2<sub>160-640</sub> CBMs, DPWs & NPFs mutated in pEGFP | This study |
| Ct-C2-GFP | *Homo sapiens* Epsin2<sub>160-640</sub> UIMs, DPWs & NPFs mutated in pEGFP | This study |
| Ct-D2-GFP | *Homo sapiens* Epsin2<sub>160-640</sub> UIMs, CBMs & NPFs mutated in pEGFP | This study |
| Ct-N2-GFP | *Homo sapiens* Epsin2<sub>160-640</sub> UIMs, CBMs & DPWs mutated in pEGFP | This study |
| Ct-ucdn2-GFP | *Homo sapiens* Epsin2<sub>160-640</sub> UIMs, CBMs, DPWs and NPFs mutated in pEGFP | This study |
| Ct-UC1N2-GFP | *Homo sapiens* Epsin2<sub>160-640</sub> DPWs & CBM2 mutated in pEGFP | This study |
| Ct-UC2N2-GFP | *Homo sapiens* Epsin2<sub>160-640</sub> CBM1 & DPWs mutated in pEGFP | This study |
| CS1 | *Homo sapiens* Epsin2<sub>160-640</sub> CBMs & DPWs mutated, CBM2 in the position of CBM1 in pEGFP | This study |
| CF1 | *Homo sapiens* Epsin2<sub>160-640</sub> CBMs & DPWs mutated, CBM2 and its flanking region in the position of CBM1 & its flanking region in pEGFP | This study |
| CS2 | *Homo sapiens* Epsin2<sub>160-640</sub> CBMS & DPWs mutated, CBM1 in the position of CBM2 in pEGFP | This study |
| CF2 | *Homo sapiens* Epsin2<sub>160-640</sub> *Homo sapiens* Epsin2<sub>160-640</sub> CBMs & DPWs mutated, CBM1 and its flanking region in the position of CBM2 & its flanking region in pEGFP | This study |
| CDN3-GFP | *Homo sapiens* Epsin3 UIM1&2 mutated in pEGFP | This study |
| UDN3-GFP | *Homo sapiens* Epsin3 CBM1 &2 mutated in pEGFP | This study |
| UCN3-GFP | *Homo sapiens* Epsin3 DPW1 to 7 mutated in pEGFP | This study |
| UCD3-GFP | *Homo sapiens* Epsin3 NPF1 to 3 mutated in pEGFP | This study |
| DN3-GFP | *Homo sapiens* Epsin3 UIMs & CBMs mutated in pEGFP | This study |
| CN3-GFP | *Homo sapiens* Epsin3 UIMs & DPWs mutated in pEGFP | This study |
| CD3-GFP | *Homo sapiens* Epsin3 UIMs & NPFs mutated in pEGFP | This study |
| UN3-GFP | *Homo sapiens* Epsin3 CBMs & DPWs mutated in pEGFP | This study |
| UD3-GFP | *Homo sapiens* Epsin3 CBMs & NPFs<br>mutated in pEGFP | This study |
| UC3-GFP | *Homo sapiens* Epsin3 DPWs and NPFs<br>mutated in pEGFP | This study |
| U3-GFP | *Homo sapiens* Epsin3 CBMs, DPWs & NPFs<br>in pEGFP mutated | This study |
| C3-GFP | *Homo sapiens* Epsin3 UIMs, DPWs & NPFs<br>mutated in pEGFP | This study |
| D3-GFP | *Homo sapiens* Epsin3 UIMs, CBMs & NPFs<br>mutated in pEGFP | This study |
| N3-GFP | *Homo sapiens* Epsin3 UIMs, CBMs &<br>DPWs mutated in pEGFP | This study |
| ucdn3-GFP | *Homo sapiens* Epsin3 UIMs, CBMs, DPWs<br>and NPFs mutated in pEGFP | This study |
| Ct-CDN3-GFP | *Homo sapiens* Epsin3<sup>176-633</sup> UIM1&2<br>mutated in pEGFP | This study |
| Ct-UDN3-GFP | *Homo sapiens* Epsin3<sup>176-633</sup> CBM1 &2<br>mutated in pEGFP | This study |
| Ct-UCN3-GFP | *Homo sapiens* Epsin3<sup>176-633</sup> DPW1 to 8<br>mutated in pEGFP | This study |
| Ct-UCD3-GFP | *Homo sapiens* Epsin3<sup>176-633</sup> NPF1 to 3<br>mutated in pEGFP | This study |
| Ct-DN3-GFP | *Homo sapiens* Epsin3<sup>176-633</sup> UIMs & CBMs<br>mutated in pEGFP | This study |
| Ct-CN3-GFP | *Homo sapiens* Epsin3<sup>176-633</sup> UIMs & DPWs<br>mutated in pEGFP | This study |
| Ct-CD3-GFP | *Homo sapiens* Epsin3<sup>176-633</sup> UIMs & NPFs<br>mutated in pEGFP | This study |
| Ct-UN3-GFP | *Homo sapiens* Epsin3<sup>176-633</sup> CBMs & DPWs<br>mutated in pEGFP | This study |
| Ct-UD3-GFP | *Homo sapiens* Epsin3<sup>176-633</sup> CBMs & NPFs<br>mutated in pEGFP | This study |
| Ct-UC3-GFP | *Homo sapiens* Epsin3<sup>176-633</sup> DPWs and NPFs<br>mutated in pEGFP | This study |
| Ct-U3-GFP | *Homo sapiens* Epsin3<sup>176-633</sup> CBMs, DPWs &<br>NPFs mutated in pEGFP | This study |
| Ct-C3-GFP | *Homo sapiens* Epsin3<sup>176-633</sup> UIMs, DPWs &<br>NPFs mutated in pEGFP | This study |
| Ct-D3-GFP | *Homo sapiens* Epsin3<sup>176-633</sup> UIMs, CBMs &<br>NPFs mutated in pEGFP | This study |
| Ct-N3-GFP | *Homo sapiens* Epsin3<sup>176-633</sup> UIMs, CBMs &<br>DPWs mutated in pEGFP | This study |
| Ct-ucdn3-GFP | *Homo sapiens* Epsin3<sup>176-633</sup> UIMs, CBMs,<br>DPWs and NPFs mutated in pEGFP | This study |
| WT3P-GFP | *Homo sapiens* Epsin3 paralog specific region mutated in pEGFP | This study |
| CDN3P-GFP | *Homo sapiens* Epsin3 paralog specific region, UIM1&2 mutated in pEGFP | This study |
| UDN3P-GFP | *Homo sapiens* Epsin3 paralog specific region, CBM1 &2 mutated in pEGFP | This study |
| UCN3P-GFP | *Homo sapiens* Epsin3 paralog specific region, DPW1 to 8 mutated in pEGFP | This study |
| UCD3P-GFP | *Homo sapiens* Epsin3 paralog specific region, NPF1 to 3 mutated in pEGFP | This study |
| DN3P-GFP | *Homo sapiens* Epsin3 paralog specific region, UIMs & CBMs mutated in pEGFP | This study |
| CN3P-GFP | *Homo sapiens* Epsin3 paralog specific region, UIMs & DPWs mutated in pEGFP | This study |
| CD3P-GFP | *Homo sapiens* Epsin3 paralog specific region, UIMs & NPFs mutated in pEGFP | This study |
| UN3P-GFP | *Homo sapiens* Epsin3 paralog specific region, CBMs & DPWs mutated in pEGFP | This study |
| UD3P-GFP | *Homo sapiens* Epsin3 paralog specific region, CBMs & NPFs mutated in pEGFP | This study |
| UC3P-GFP | *Homo sapiens* Epsin3 paralog specific region, DPWs and NPFs mutated in pEGFP | This study |

